# The polyamine naphthyl-acetyl spermine trihydrochloride (NASPM) lacks specificity for Ca^2+^-permeable AMPA receptors and suppresses seizure like activity in human brain tissue by inhibition of NMDA receptors

**DOI:** 10.1101/2025.05.14.653650

**Authors:** Alice Podestà, Laura Monni, Franziska Arnold, Julia Arnold, Sara Bertelli, Ran Xu, Julia Onken, Ulrich-Wilhelm Thomale, Thilo Kalbhenn, Matthias Simon, Thomas Sauvigny, Henrik Alle, Andrew J. R. Plested, Martin Holtkamp, Jörg R.P. Geiger, Pawel Fidzinski

## Abstract

For decades, naphthyl-acetyl spermine trihydrochloride (NASPM) has been used as a selective inhibitor of calcium-permeable AMPA receptors (CP-AMPAR). In rodents, NASPM is known to suppress seizures *in vivo* and seizure-like events (SLE) *in vitro*, suggesting possible involvement of CP-AMPAR in ictogenesis and epileptogenesis. To address whether these findings can be translated to human brain, we investigated the involvement of glutamatergic receptor subclasses in SLE in human cortex *ex vivo,* demonstrating that glutamatergic receptor antagonists can block (NASPM and APV) or reduce (UBP302, GYKI52466, GYKI53655) SLE. Using a multimethodological approach we were able to demonstrate that both NASPM and APV inhibit human SLE by inhibition of NMDA receptors. Our results further show that the inhibitory effect of NASPM on NMDA receptors is sufficient to explain its inhibition of seizure like activity, rather than its action on CP-AMPA receptors. Thus, our findings challenge previous knowledge on the use of NASPM as a specific CP-AMPAR inhibitor. Some phenomena previously attributed to CP-AMPAR, may need to be re-examined more closely. Overall, our study raises awareness about potential pitfalls in the use of existing pharmacological agents and sets a new paradigm for the use of NASPM in neuroscience research while questioning its therapeutic potential in a clinical context.

## Introduction

Epilepsy is a chronic neurological disorder characterised by increased propensity for spontaneous seizures. Functionally, epileptic seizures are attributed to synchronous discharges of neuronal populations caused by increased intrinsic activity which can result from an imbalance between excitation and inhibition within the neuronal network. Aberrant activity of ionotropic and metabotropic glutamatergic receptors is strongly correlated with epilepsy and nevertheless there remains limited understanding how and at which level glutamatergic receptor subtypes are involved in ictogenesis and epileptogenesis (Hanson et al., 2024; Hanada et al., 2020; Bonansco et al., 2016; Rogawski et al., 2013; Ghasemi et al., 2011; Rogawski *et al.,* 1992). This knowledge gap is particularly pronounced in the context of human tissue.

Vertebrate ionotropic glutamate receptors can be classified into four subcategories: alpha-Amino-3-hydroxy-5-methylisoxazole-4-propionate receptors (AMPAR), N-methyl-D-aspartate receptors (NMDAR), kainate receptors (KAR) and the delta subtype. AMPAR are widely expressed in central nervous system and consist of tetrameric receptors formed by the combination of four core subunits (GluA1-4). The receptor composition varies across brain regions and during development; generally complexes containg GluA1, GluA2 and GluA3 are the most abundant (Hanson et al., 2024; Zhao et al., 2019; Shepherd et Huganir, 2007; Kumar et al., 2002). Among AMPAR subunits, GluA2 undergoes RNA editing which determines the expression of an arginine residue in the pore region and a consequent low permeability to divalent cations. Conversely, GluA2 lacking AMPAR or AMPAR with unedited GluA2 subunits are permeable to divalent cations, specifically calcium (Greger et al. 2017; Shepherd and Huganir, 2007; Burnashev et al. 1992; Sommer et al. 1991; Boulter et al., 1990). In rodents, most GluA2 transcripts in the adult mammalian brain undergo editing, therefore resulting in predominance of AMPAR with a low calcium permeability (Kawahara et al., 2005; Isaac et al. 2007). In contrast, Ca^2+^-permeable AMPAR (CP-AMPAR) are preferentially found in GABAergic interneurons (Lalanne et al., 2018; Lalanne et al., 2016; Geiger et al., 1995; Koh et al., 1995a; Jonas et al., 1994; McBain and Dingledine, 1993). Abnormal expression of CP-AMPAR has been found to be involved in various neurodegenerative diseases, ischemia and epilepsy, likely contributing to elevated Ca^2+^ influx and consecutive excitotoxicity (Guo et Ma, 2021; Salpietro et al., 2019; Kawahara et al. 2007). The link between the GluA2 editing and epilepsy has been demonstrated in animal models suggesting a positive association between seizure propensity and CP-AMPAR expression levels. It is unknown, however, whether the same association could apply to the human brain (Konen et al., 2020; Vollmar et al., 2004; Kortenbruck et al., 2001; Higuchi et al., 2000; Grigorenko et al., 1998; Brusa et al., 1995).

The functional impact of CP-AMPAR in ictogenesis has been investigated in animal models using selective inhibitors of CP-AMPAR activity, such as the Joro spider toxin JSTX, the synthetic compound IEM-1460 and the polyamine drug naphthyl-acetyl spermine trihydrochloride (NASPM).

Since the initial description by Kanai et al. which reported the anti-seizure effect of NASPM *in vivo*, this compound has been widely used for investigating CP-AMPAR function, mechanism and activity (Kanai et al., 1992). Consequently, several studies have used NASPM to further investigate its anti-seizure properties related to CP-AMPAR inhibition (Twomey et al., 2018; Tóth and McBain, 1998; Washburn et al., 1997; Washburn and Dingledine, 1996; Bowie and Mayer, 1995; Blaschke et al., 1993; Herlitze et al., 1993; Kanai et al., 1992). Whether CP-AMPAR are involved in human seizures is not known. Current literature is based on animal models, and the translation of these findings to humans might not be reliable. To address this gap, we investigated the involvement of ionotropic glutamatergic receptor subclasses in seizure-like activity in an *ex vivo* human brain model. We employed a pharmacological approach based on putative subunit and/or subclass-specific inhibitors. We were able to demonstrate that nearly all classes of glutamatergic receptor contribute to seizure- like activity in human brain. Most importantly, we found that the putative CP-AMPAR-selective drug NASPM fully blocks seizure-like activity in human brain but does so through inhibition of NMDA receptors. These results bring into question the previous use of NASPM as a drug specifically affecting CP-AMPAR.

## Material and Methods

### Resection of human brain samples

Surgical specimens were obtained from the Neurosurgery Department of Charité Universitätsmedizin Berlin (30 patients), Neurosurgery Department Bethel Klinikum Bielefeld (35 patients); Neurosurgery Department of Universitätsklinikum Hamburg-Eppendorf, Hamburg (12 patients). All patients provided informed consent for tissue donation and the study was positively assessed by the local ethical committee (Berlin: vote no. EA2/111/14, ethical commission of the Charité, Bielefeld: vote no.2020-517-f-S, ethical commission of the medical chamber Westfalen Lippe, Hamburg: vote no. 2023-200674-BO-bet, ethical commission of the medical chamber Hamburg). All experiments were performed in accordance with the relevant guidelines and regulations. Tissue was collected from 77 patients 73.17% of which resulted in viable tissue. The ages of the patients ranged from 0 to 75 years with no apparent difference between viable and non- viable material along the age distribution (**Figure 1, B**). Resections were performed with clean cuts along the edges of the resected tissue, except when such cuts contradicted therapeutic requirements. Scalpel incisions were perpendicular to the pial surface aiming to prevent abrupt disruption of neuronal morphology throughout the cortical depth. Each surgically resected specimen was immediately placed in a sterile container filled with sterile high saccharose-based artificial cerebrospinal fluid (aCSF) composed of (in mM): 2.5 KCl, 1.25 NaH_2_PO_4_, 25 NaHCO_3_, 10 glucose, 0.5 CaCl_2_·4H_2_O and 3 MgCl_2_, 75 Saccharose with the osmolality of 310–320 mOsm/kg. The transport solution was pre-chilled to 2–4 °C and thoroughly bubbled with carbogen (95% O_2_/5% CO_2_) gas prior to collection. The tissue was transported from operation theatre to the *in situ* remote laboratory in gas-tight bottles to maintain pH and O_2_ levels stable. Transport time from the operating theatre to the laboratory was between 15–30 min. Neocortical resections had a typical volume of ∼3-5 cm³, yielding up to 15-30 viable slices for electrophysiological experiments which were distributed between various experimenters and projects. Sliced samples were transported to Berlin laboratory either directly or through a custom-made transport system to preserve slice quality. Total transport time ranged from 15 min to 5 hours without noticeable differences in tissue quality (**Figure 1, C-D**). Slices were incubated for at least 2 hours in high saccharose-based artificial cerebrospinal fluid (aCSF) before performing experiments **(Figure 1, D).**

**Figure 1.**
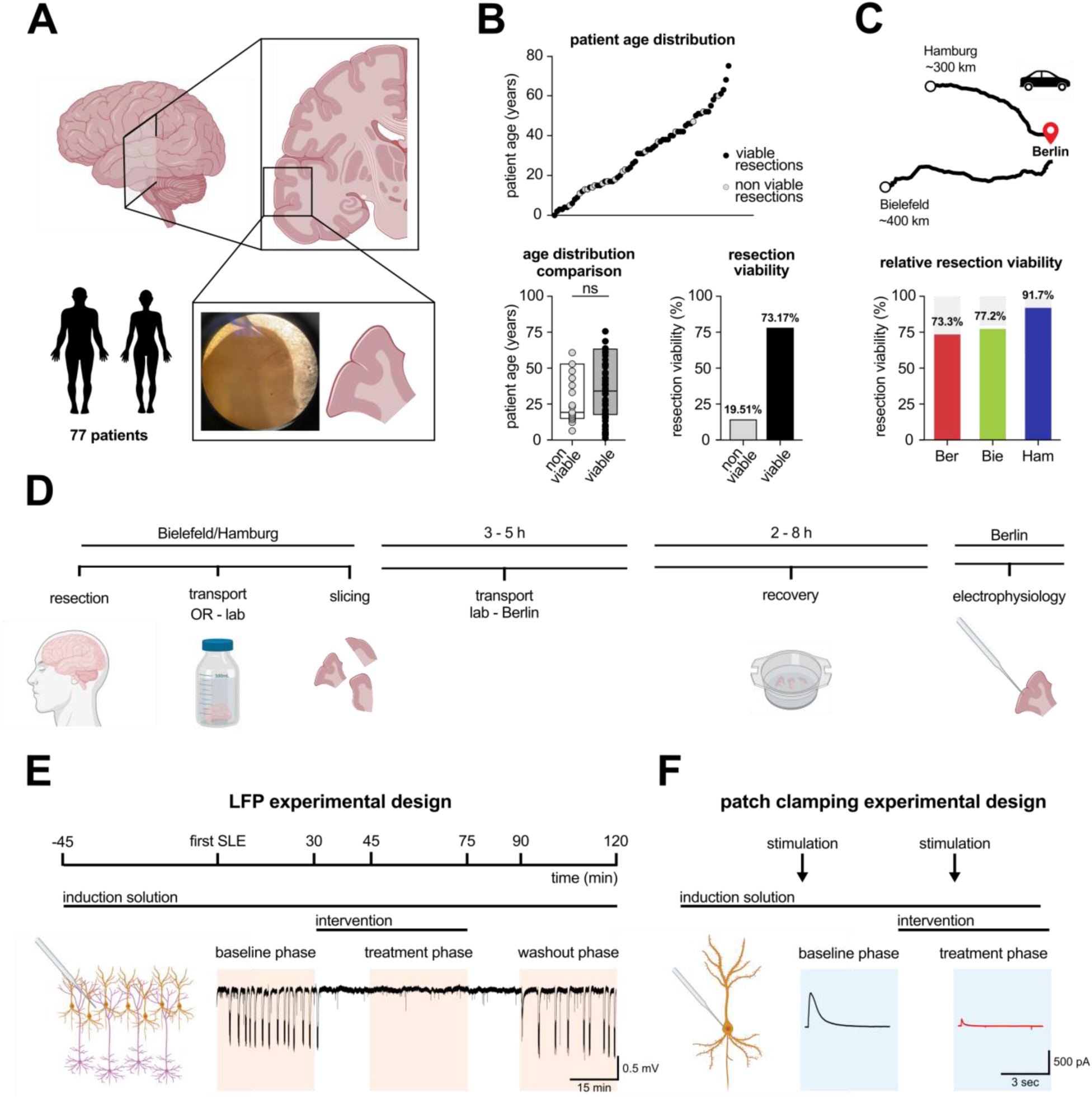
Experimental setting for human tissue experiments. **A.** Schematic representation of area of resection. Brain samples were obtained from male and female patients aged between 0 to 75 years. **B.** Age distribution of patients. Grey dots represent resection with low or no viability (19.51% of resections), black dots represent viable resections (73.17%). Age distribution for viable (black) and not viable (grey) resection is shown as scatter plot; superimposed box plots show the median, and interquartile range. Percentage of viability of the resections is shown as bar chart. <u>Statistics:</u> Mann-Whitney test; normality assessed via Shapiro-Wilk test. **C.** Resected material was supplied from three different centres located in Berlin, Bielefeld and Hamburg. Relative viability for single centre is shown as column chart. **D.** Schematic overview of the experimental workflow, from tissue resection to electrophysiological recordings. **E.** LFP experimental setup. Schematic timeline of SLE induction and pharmacological intervention. A representative 0Mg-GBZ-CGP recording trace is shown as a reference, to improve visualization, slow drift artifacts have been removed. **F.** Patch-clamp experimental setup. Whole-cell voltage-clamp recordings were performed during SLE induction and pharmacological intervention. Evoked EPSCs were induced by electrical stimulation at the beginning of each phase. Representative EPSC traces from different phases are shown for reference. To enhance clarity, 50 Hz noise has been removed.

### Human neocortical slice preparation

Tissue was visually inspected for mechanical or thermal damage and to identify the most suitable orientation for slice preparation. Meninges were carefully removed using blunt forceps. Specimen were cut to planar surface and glued perpendicularly to the pial surface to ensure an optimised cutting angle for preservation of neurite integrity. Tissue was cut with a Leica VT 12000S vibratome at the speed of 0.07 mm/min, oscillation rate was dependent on the presence of residual meninges and was set to the amplitude of 1-2 mm with a cutting frequency of 80 Hz. Specimen were sliced at the thickness of 450 µm in chilled sterile high saccharose-based aCSF with continuous carbogenation (95% O_2_/5% CO_2_). Slices were transferred to a custom-made sterile interface system filled with pre-warmed (35°C) high saccharose-based aCSF for 30 min. After warm-recovery, slices were incubated for at least additional 2h at room temperature. Upon completion of total recovery period, slices were stored in a sterile interface system at room temperature for the total duration of the experimental time which lasted between 6 and 72h.

### Heterologous expression of NMDA receptors

Experiments were conducted using ND7/23 cells (Wood et al., 1990). Cells were cultivated in Dulbecco’s Modified Eagle Medium (DMEM), supplemented with 6% FBS, and maintained at 37°C and 5% CO_2_ in a humidified incubator. They were passaged twice a week and seeded onto glass coverslips in 60mm Petri dishes two days prior to experiments.

pCI-EGFP-NR2a wt (Addgene plasmid #45445, http://n2t.net/addgene:45445; RRID:Addgene_45445) and pCI-EGFP-NR1 wt (Addgene plasmid #45446, http://n2t.net/addgene:45446; RRID:Addgene_45446) were gifts from Andres Barria & Robert Malinow (Barria et Malinow, 2002). The NR2a cassette was subcloned in pcDNA3.1 and EGFP was replaced by mScarlet. NR1 was subcloned into pcDNA 3.1 (without the EGFP). ND7/23 cells were transiently co-trasfected with 2 µg of a 1:1 plasmid mixture and polyethylenimine (PEI) in a 1:3 volume ratio. Cells were selected for experiments based on red fluorescence signals.

Micropipettes with a tip resistance of 2-3 MΩ were prepared from borosilicate glass using a Sutter P-1000 micropipette puller and filled with intracellular solution containing 120 mM CsCl, 10 mM CsF, 10 mM NaCl and 2 mM EGTA adjusted to 265 mOsm/L and pH 7.35. Coverslips were transferred to the bath mounted on an Olympus IX-81 microscope 16-48 h after transfection and bathed in extracellular solution (145 mM NaCl, 10 mM HEPES, 10 mM Glucose, 2.5 mM KCl and 2 mM CaCl

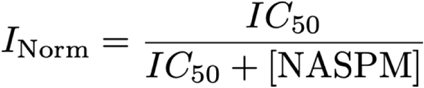

pH 7.35, 295 mOsm/L). Whole-cell currents were recorded using an AxoPatch 200B amplifier (Molecular Devices) and the AxoGraph acquisition software (Axograph Scientific). Single lifted cells were perfused with standard extracellular solution or solutions containing either 87 µM, 26.1 µM, 8.7 µM of 1-Naphthylacetyl spermine. NASPM was added the day of the experiment. NMDA receptor responses were evoked by 100 µM NMDA and 100 µM glycine using rapid solution exchange, achieved using custom-made glass perfusion tools (Plested et al., 2021) mounted on a piezoelectric actuator (PICMA P-840.40, Physik Instrument, Karlsruhe, Germany). The extracellular solution used for fast perfusion was pre-warmed (37°C) for a mild degassing, and then allowed to cool before use. Experiments were performed at room temperature.

Data were analyzed using IgorPro and the NeuroMatic plug-in (WaveMetrics). NASPM currents were normalized as the ratio of the peak response to the NMDA-evoked response immediately preceding co-application. To estimate *IC*_50_ values, we performed fits to the three sets of normalized peak currents at different voltages, using a Langmuir isotherm (assuming a single binding site for NASPM):

### Extracellular recordings

Human slices were transferred to an interface chamber and continuously perfused with carbogenated aCSF solution at flow rate of 2mL*min^-1^. aCSF was composed of (in mM): 129 NaCl, 1.25 NaH_2_PO_4_, 1.6 CaCl_2_, 3 KCl, 1.8 MgSO_4_, 21 NaHCO_3_, 10 C_6_H_12_O_6_ if not otherwise specified. Extracellular local field potential (LFP) recordings were conducted using borosilicate pipettes (Science Products, Hofheim, Germany; 1.5 mm outer diameter) of electrical resistance of 1-2 MΩ filled with 150 mM NaCl. Signals were acquired from neocortex layer II with custom-made amplifiers (10×) connected to an AD converter (Micro 1401 mk II, Cambridge Electronic Design Limited, Cambridge, United Kingdom). Data were recorded using Spike2 and Signal (versions 7.00 and 3.07, respectively, Cambridge Electronic Design Limited, Cambridge, United Kingdom). Seizure like events (SLE) in human cortical slices were induced by applying various aCSF-based pharmacological induction methods at controlled osmolarity and temperature (35°C, 305 ± 5 mOsm/L, pH 7.4). The induction methods used were the following:

i. High K^+^ aCSF + 100 µM 4-AP. Composition (in mM) 122 NaCl, 1.25 NaH_2_PO_4_, 1.6 CaCl_2_, 10 KCl, 1.8 MgSO_4_, 21 NaHCO_3_, 10 C_6_H_12_O_6_ referred as HighK+4AP;
ii. 0 Mg^2+^ aCSF+ 10 µM bicuculline (BIC). Composition (in mM): 130.8 NaCl, 1.25 NaH_2_PO_4_, 1.6 CaCl_2_, 3 KCl, 21 NaHCO_3_, 10 C_6_H_12_O_6_ referred as 0Mg+BIC;
iii. 0 Mg^2+^ aCSF + 10 µM gabazine (GBZ) + 10 µM CGP 55845. Composition (in mM) 130.8 NaCl, 1.25 NaH_2_PO_4_, 1.6 CaCl_2_, 3 KCl, 21 NaHCO_3_, 10 C_6_H_12_O_6_, referred as 0Mg+GBZ+CGP;

SLE in mouse tissue were induced by applying the following aCSF-based induction at controlled osmolarity and temperature (35°C, 305 ± 5 mOsm/L, pH 7.4):

a. 0 Mg^2+^ aCSF. Composition (in mM) 130.8 NaCl, 1.25 NaH_2_PO_4_, 1.6 CaCl_2_, 3 KCl, 21 NaHCO_3_, 10 glucose;
b. _(b)_ aCSF + 100 µM 4-AP. Composition (in mM) 129 NaCl, 1.25 NaH_2_PO_4_, 1.6 CaCl_2_, 3 KCl, 1.8 MgSO_4_, 21 NaHCO_3_, 10 C_6_H_12_O_6;_
c. _(c)_ aCSF + 10 µM bicuculline (BIC) + 10 µM CGP 55845. Composition (in mM) 129 NaCl, 1.25 NaH_2_PO_4_, 1.6 CaCl_2_, 3 KCl, 1.8 MgSO_4_, 21 NaHCO_3_, 10 C_6_H_12_O_6;_

To assess the involvement of glutamatergic receptors in SLE, a combination of glutamatergic receptor antagonists was added. Specifically, following drugs were used: 12.5 µM UBP302, a selective kainate receptor (KAR) antagonist; 50 µM GYKI53655, a selective AMPA receptor (AMPAR) antagonist; 30 µM GYKI52466, an antagonist for KAR and AMPAR; or a combination of these (50 µM GYKI53455 + 12.5 µM UBP302) together with 10 µM NBQX, a non-selective AMPAR antagonist. To investigate NMDAR involvement, 50 µM APV was used (**Figure1, E**).

### Intracellular recordings from human cortical neurons

Human slices were transferred to a submerged high flow rate (10 mL*min^-1^) chamber and perfused with carbogenated aCSF solution at monitored temperature and osmolarity (35°C, 305 ± 5 mOsm/L, pH 7.4). Borosilicate pipettes (Science Products, Hofheim, Germany; 1.5 mm outer diameter) with an electrode resistance 4-5 MΩ were pulled with a vertical puller (PC-10, Narishige, Tokyo, Japan). Patch-clamp somatic recordings in whole-cell configuration were performed in acute human slices from neocortical layer II-III by using borosilicate pipettes filled with Cs-based internal solution of composition (in mM): 130 CsCl, 2 MgCl_2_, 10 HEPES, 2 MgATP, 0.2 EGTA, 5 QX-314, 0.2% biocytin, osmolarity 290 ± 5 mOsm/L, pH 7.2-7.4. Liquid junction potential was calculated to be +4 mV.

### NMDAR mediated currents in ex vivo human brain slices

NMDAR mediated excitatory postsynaptic currents (NMDAR-EPSC) were recorded in voltage-clamp configuration the membrane potential of +40 mV using 0Mg+GBZ+CGP as extracellular solution. To isolate NMDA currents, a combination of pharmacological blockers was used to block AMPA receptors, kainate receptors and GABA-A receptors (12.5 µM UBP302; 50 µM GYKI53655; 10µM NBQX, and gabazine + CGP as stated above) and the NMDAR co-agonist glycine (10µM) was applied continuously. NMDAR-eEPSCs were evoked by extracellular electrical stimulation (duration 100µs) using a borosilicate pipette filled with 150 mM NaCl (electrical resistance of 1-2 MΩ). To assess drug effects on NMDAR-EPSCs, 100 µM NASPM, 50 µM APV or milli-Q water as vehicle were used (**Figure 1, F**). Access resistance (*R*_a_) and series resistance (*R*_s_) were monitored during recordings to ensure their stability over time. Quality criteria for recording inclusion were (i) a cell-attached seal > 1 GΩ before establishing whole cell configuration; (ii) stable series resistance < 30 MΩ with and (iii) a *R*_s_/*R*_a_ ratio > 3. Cell resting membrane potential was measured within the first minute after establishing whole-cell configuration and used as indication of functional integrity. Cell with resting potential < -70 mV or > -50 mV were excluded.

### Data analysis

SLE were identified by the following criteria: (i) depolarisation of the field potential of > 0.5 mV (ii) for > 10 s, and (iii) superimposition of ripple-like discharges. Slices were monitored for presence of SLE for 45 min after perfusion of the induction solution; reactivity to seizure like event induction varied across tested induction methods, slices which did not exhibit stable seizure like activity within this time were excluded, resulting in 46.53% of excluded slices. After occurrence of the first SLE, a 30 min baseline was established, after which the drugs or vehicle solution were perfused and SLE activity was monitored for a further 45 min during this treatment phase. After this time, the drugs were washed out and SLE activity was monitored for additional 45 min. For analysis, the final 30 minutes of each phase (baseline, treatment, and washout) were used (**Figure 1, E**).

Electrophysiological recordings were processed and analysed with custom-written Matlab code (R2015b-2024, MathWorks, Natick, MA, United States, code available upon request). Representative trace and figure processing were performed with GraphPad Prism 10, Matlab and Inkscape 1.1 (The Inkscape Project).

### Statistical analysis

In each experimental group, tissue from at least 5 patients was used. The number was adjusted to find appropriate balance between tissue availability and adequate sample size with regard to parameter variability. Data were analysed with GraphPad Prism 10 (GraphPad Software Inc., San Diego, CA, USA). Values of p < 0.05 were considered as statistically significant. Differences in SLE incidence across various conditions were evaluated using one-way ANOVA with Dunnett’s T3 multiple comparison test. Significant differences of SLE properties were calculated with Kruskal-Wallis test with Dunn’s multiple comparison test. In intervention experiments, SLE incidence, amplitude, duration, spike frequency within SLE and spike number were evaluated in within-condition (Baseline vs. Treatment, Baseline vs. Washout) using one-way ANOVA or Friedman test with Dunn’s multiple comparison test where appropriate. In case of full SLE block during pharmacological intervention only incidence rate was taken into consideration. For assessment of glutamatergic involvement in SLE incidence and properties, 0Mg+GBZ+CGP protocol was used as reference method. Kruskal-Wallis with Dunn’s multiple comparisons was used to determine statistical differences across protocols. In silico assessed binding affinity of NASPM for NMDAR, CP-AMPAR and AMPAR was compared across the condition via Kruskal-Wallis test with Dunn’s multiple comparisons test. Normality distribution of the data was assessed with Shapiro-Wilk test.

### Materials

4-aminopyridine (A54008), DMSO (D1435) were purchased from Sigma, Munich, Germany. APV (ab120271) was purchased from Abcam Limited, Cambridge, UK. NASPM trihydrochloride (2766, *ex vivo* experiments), CGP 55845 (1248), UBP 302 (2079), GYKI52466 (1454), GYKI53655 (2555), QX 314 chloride (2313), NBQX (373) were purchased from Tocris, Bristol, UK. Gabazine (HB0901) and NASPM (HB0441, heterologous expression experiments) was purchased from Hello Bio, Bristol, UK.

## Results

### Induction of seizure like events in *ex vivo* human neocortical slices

To investigate SLE in human brain tissue, we used three different induction methods in cortical slices from 40 patients with an age range of 0-68 years. SLE were induced by using i) “HighK+4AP”: aCSF with elevated potassium (10 mM K^+^) coupled with potassium channel block by 4-aminopyridine (4AP); ii) “0Mg+BIC”: aCSF with magnesium omitted (nominal 0 mM Mg^2+^) together with GABA_A_ receptor block by bicuculline; iii) “0Mg+GBZ+CGP”: aCSF with 0 mM Mg^2+^ together with GABA_A_ receptor block by gabazine and GABA_B_ receptor block by CGP55845. In human cortical slices, the success rate of induction was different across the three methods: HighK+4AP induction resulted in SLE in only 25% of exposed slices while 56.3% of slices where 0Mg+GBZ+CGP was used showed robust SLE activity (**Figure 2, B**). When SLE activity was successfully induced, the SLE were stable and mostly lasted > 3 h. 0 Mg^2+^ aCSF based induced SLE with similar electrophysiological pattern and features (0Mg+BIC: 0.43 ±0.18 events/min. 37 slices from 11 patients; 0Mg+GBZ+CGP: 0.40 ±0.18 events/min, 40 slices from 16 patients; **Figure 2A-B**), while HighK-4AP induction method induced fewer but longer SLE (incidence: 0.29 ± 0.24 events/min, p= 0.0235, 21 slices from 12 patients; duration: 135.81±0.18 sec; p=0.0001, p=0.0196, 14 slices from 10 patients; **Figure 2A-B**). While spike numbers remained consistent across all induction protocols, HighK+4AP-induced SLE showed a notably lower spike frequency due to their extended duration (p=0.0051, p=0.0007, 14 slices from 10 patients; **Figure 2A-B**).

**Figure 2.**
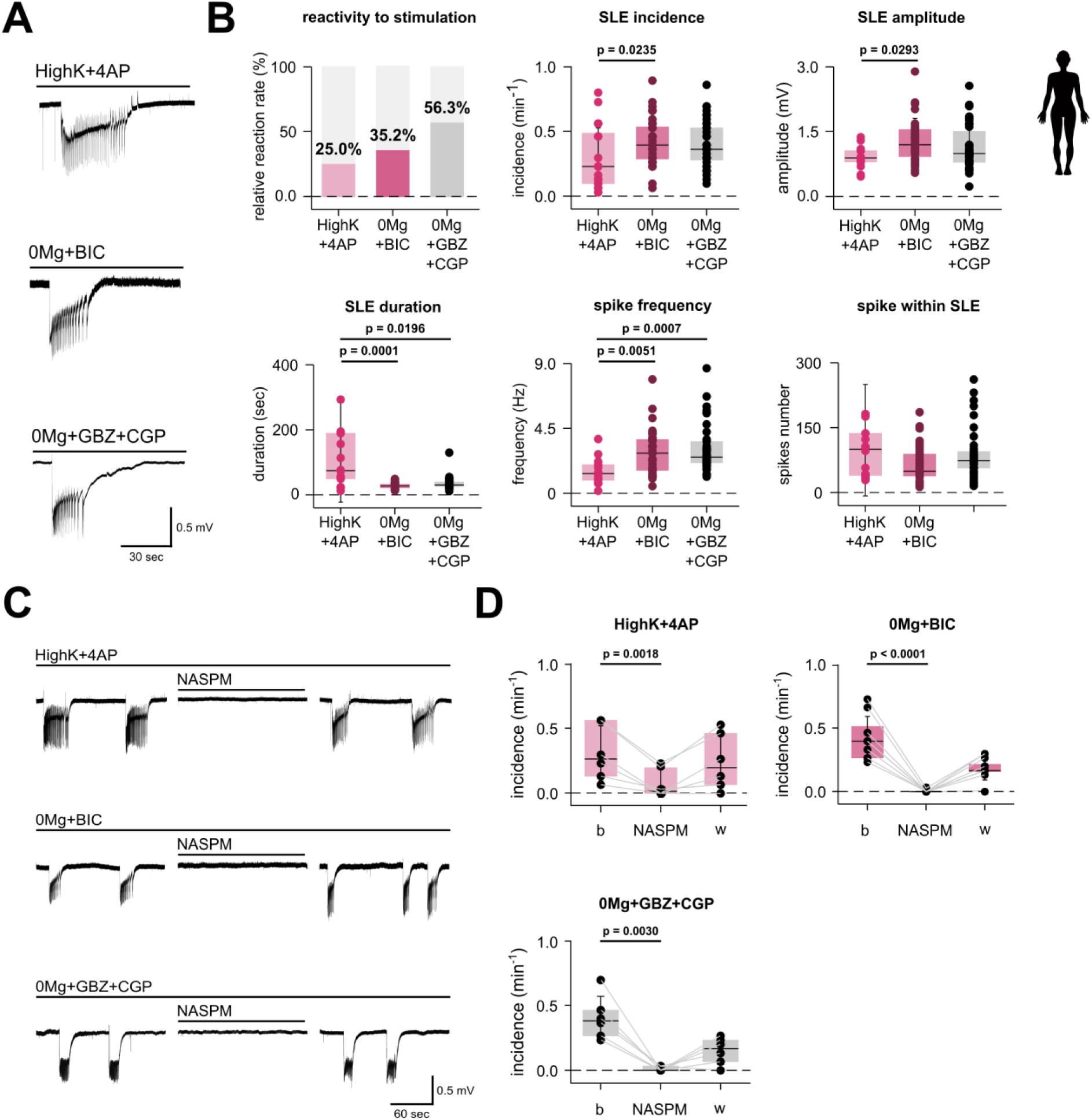
SLE induction in human brain cortex and NASPM action on SLE incidence. **A.** Representative seizure like events induced by three different induction methods. Extracellular recordings from layer II/III of human temporal cortex in different experimental settings. **B.** Comparison of SLE incidence and properties across induction methods. HighK+4AP induction method produce SLE with significant lower incidence (p = 0.0235) and amplitude (p = 0.0293) compared to 0Mg+BIC induction method. HighK+4AP evoke significantly longer SLE (p= 0.0001; p = 0.0196) without affecting spike number, thus having a significant lower spike frequency within single SLE when compared to other induction methods (p=0.0051; p = 0.0007). <u>Statistics:</u> Kruskal-Wallis test with Dunn’s multiple comparison test. Normality was calculated with Shapiro-Wilk test. Sample size: HighK-4AP: n=21 slices from 12 patients; 0Mg+BIC : n=37 slices from 11 patients, 0Mg+GBZ+CGP: 40 slices from 16 patients. **C.** Applying 100 µM NASPM completely but reversibly inhibits SLEs, independent of induction method. For each induction method a representative portion of baseline, NASPM intervention and washout phase are shown, slow baseline drift of the traces was removed for visualization. **D.** Effect of 100 µM NASPM on SLE incidence across the various methods during baseline (b), treatment (NASPM) and washout (w) phase. NASPM intervention significantly reduce SLE incidence when induced by HighK+4AP (p= 0.0018, n=6 slices from 5 patients); 0Mg+BIC (p<0.0001, n=9 slices from 5 patients) and 0Mg+GBZ+CGP (p= 0.0030, n=6 slices from 6 patients). The effect is partially restored upon washout (w). Data are shown as scatter plot of individual recording; the superimposed box plots show the median (centre line), interquartile range (box edges), and standard deviation (whiskers). <u>Statistics</u>: Friedman test with Dunn’s multiple comparison test. Normality was calculated with Shapiro-Wilk test.

### Human seizure like events are inhibited by NASPM but not IEM-1460

To investigate the involvement of CP-AMPAR in human ictogenesis, we tested the effects of NASPM and IEM-1460 on human SLE. In slices in which SLE were induced by HighK+4AP, NASPM significantly reduced SLE activity (HighK+4AP: p=0.0018, 6 slices from 6 patients, **Figure 2, C-D**). In animal models, CP-AMPAR are known to be preferentially expressed in fast-spiking GABAergic interneurons (Lalanne et al., 2018; Lalanne et al., 2016). We therefore hypothesized that in the presence of GABAergic blockers when using 0Mg+BIC or 0Mg+GBZ+CGP induction methods, CP- AMPAR inhibition should have no or little effect on SLE activity. To our surprise, 100 µM NASPM also strongly inhibited SLE activity when using 0Mg+BIC or 0Mg+GBZ+CGP as induction (0Mg-BIC: p<0.0001, 9 slices from 6 patients; 0Mg+GBZ+ CGP: p=0.0030, 6 slices from 6 patients, **Figure 2, C-D**). In contrast, application of 150 µM IEM-1460, another CP-AMPAR inhibitor based on polyamine toxins, did not alter SLE incidence (0Mg+BIC: p>0.999; 9 slices from 7 patients, HighK+4AP: p>0.999; 3 slices from 3 patients **Supplementary Figure 1, A-B**), suggesting different mechanisms of action (and targets) for NASPM and IEM-1460 in human brain tissue.

### Glutamatergic receptors are involved in seizure like events in human cortical slices

The divergent effects of NASPM and IEM-1460 on human SLE activity prompted us to investigate the involvement of glutamatergic receptors in more detail. To this end, we induced SLE in human brain slices (0Mg+GBZ+CGP) in the presence of AMPA and KA receptor inhibitors either selectively or in combination: kainate receptors were blocked by 12.5 µM UBP302; AMPA receptors were blocked by 50 µM GYKI53655 and a combined block of kainate and AMPA receptors was established by 30 µM GYKI52466 or by 50 µM GYKI53655 + 12.5 µM UBP302 + 10 µM NBQX; **Figure 3**). None of these manipulations prevented SLE induction, although they resulted in a general trend of reduced SLE incidence (0Mg+GBZ+CGP + GYKI52466: p=0.0059, 17 slices from 7 patients; 0Mg+GBZ+CGP + GYKI53655+UBP302+NBQX: p=0.0311, 16 slices from 10 patients, **Figure 4, A**), suggesting that the inhibited receptor subclasses are necessary for SLE maintenance. Spike frequency, and to a lesser extent spike number, were both reduced when AMPAR were inhibited, suggesting AMPAR involvement in the generation of ictal population spikes (spike frequency: 0Mg+GBZ+CGP +GYKI53455: p<0.0001, 11 slices from 7 patients; 0Mg+GBZ+CGP +GYKI52466: p=0.0001, 17 slices from 7 patients; 0Mg+GBZ+CGP +GYKI53655+UBP302+NBQX: p=0.0002, 16 slices from 10 patients. Spike number: 0Mg+GBZ+CGP +GYKI53655: p=0.0024, 11 slices from 7 patients; 0Mg+GBZ+CGP +GYKI52466: p=0.0017, 17 slices from 7 patients, **Figure 4, A-B**). SLE amplitude and duration were not affected by AMPA and kainate blockers (**Figure 4, A-B; Supplementary Figure 1, C**). In the 0Mg+GBZ+CGP induction method and in presence of 30 µM GYKI52466 (p=0.0144, 17 slices from 7 patients) or 50 µM GYKI53655 + 12.5 µM UBP302 + 10 µM NBQX, 100µM (p=0.0248, 16 slices from 10 patients) NASPM abolished SLE incidence during the treatment phase. In the presence of GYKI53655 alone, NASPM showed a strong decreasing trend and reached significance during washout phase (p=0.0228, 11 slices from 7 patients). In presence of 12.5 µM UBP302, the effect of 100µM NASPM was less than for the other conditions, but the baseline rate of SLE was also mildly reduced, consistent with partial involvement of kainate receptors in human SLE (**Figure 4, C**).

**Figure 3.**
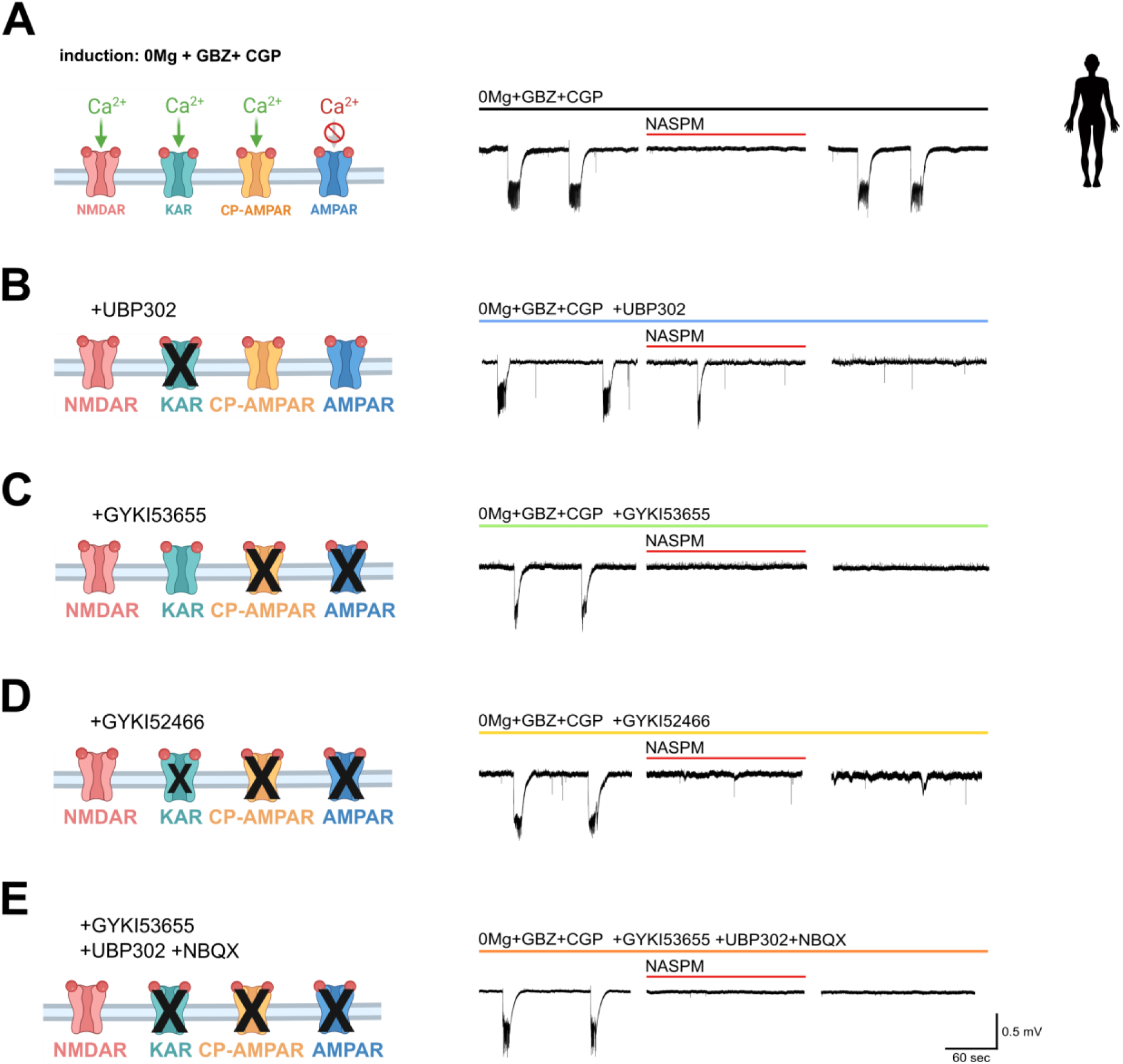
Schematic block of various glutamatergic receptor and relative representative traces. Schematic representation of inhibited receptor families for each drug (left), representative portion (300 sec) of each experimental phases (baseline, treatment and washout) it’s shown for each relative induction method (right). For experimental setting see Figure 1. SLE are induced by 0Mg+GBZ+CGP (control, **A**) in the presence of various glutamatergic blockers: the KAR selective antagonist UBP302 (**B**), AMPAR antagonists GYKI53655 (**C**); the AMPAR antagonist GYKI52466 which, to a lower extent, also inhibity KAR (**D**), and a combination of UBP302, GYKI53655, and NBQX to block simultaneously AMPARs and KAR (**E**).

**Figure 4.**
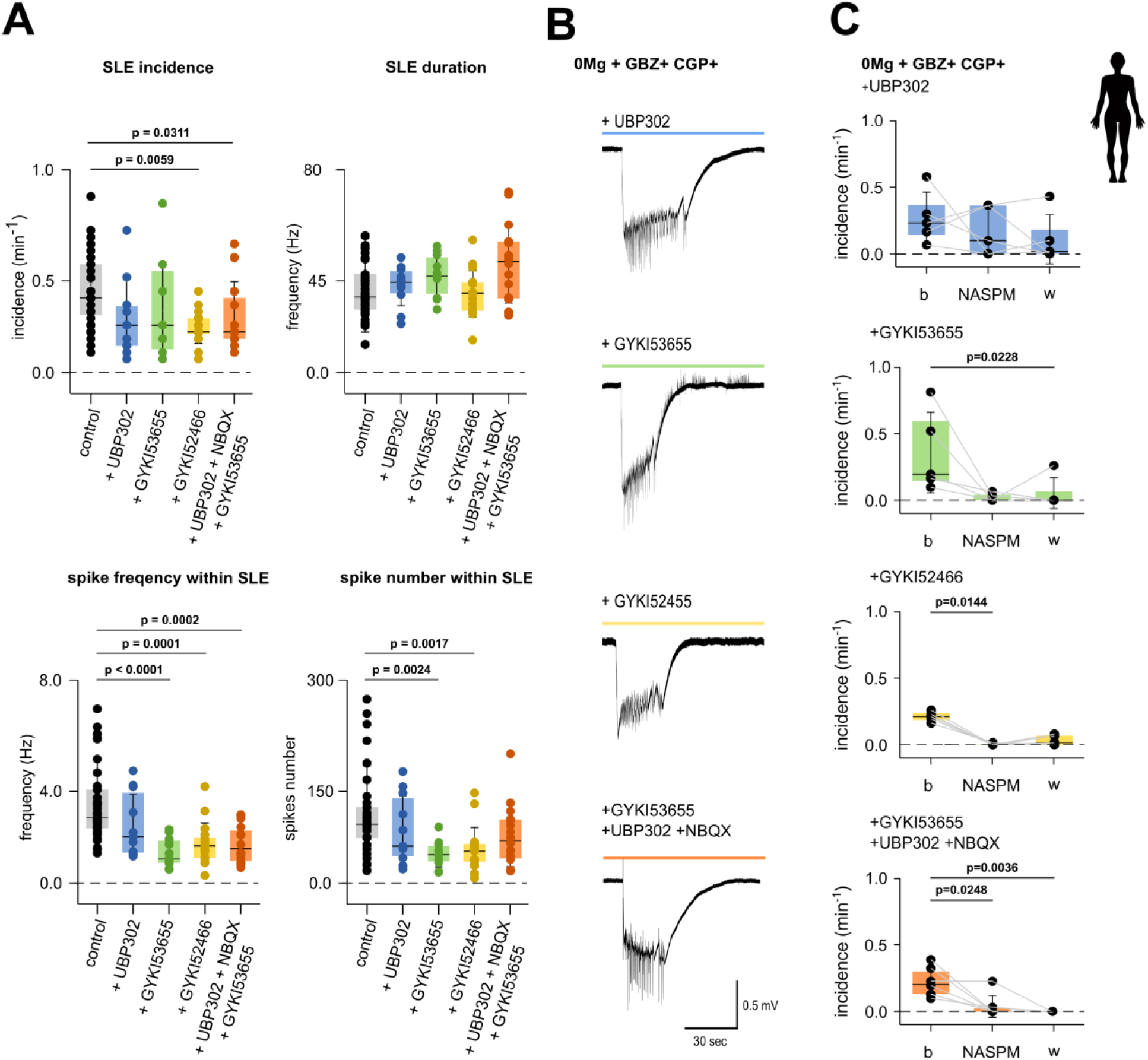
Properties and incidence of SLEs vary in presence of various glutamatergic receptor antagonists. **A.** SLE incidence and properties are influenced by glutamatergic receptors antagonists. SLE incidence significantly decreased in presence of a combined inhibition of kainate and AMPA receptors using either GYKI52466 (p= 0.0059) or the combination of GYKI53655 (50 µM), UBP302 (12.5 µM), and NBQX (10 µM, p=0.0311). Spike frequency within SLEs was likewise reduced by GYKI53655 (50 µM, p<0.0001), by GYKI52466 (30 µM, p=0.0001), and by the combined block of 12.5 µM UBP302, 50 µM GYKI53655 and 10 µM NBQX (p = 0.0002). Similarly, spike number within SLE was reduced in presence of 50µM GYKI53655 (p=0.0024) or 30µM GYKI52466 (p=0.0017). **B.** Representative seizure like events induced by 0Mg+GBZ+CGP in presence of the glutamatergic receptor blockers investigated. **C.** Effect of NASPM (100 µM) had no major effect on SLE incidence in the presence of 12.5 µM UBP302. However, it completely abolished SLEs occurring in presence of GYKI52466 (30 µM, p=0.0144) or with the combined blockade of 12.5 µM UBP302, 50 µM GYKI53655 and 10 µM NBQX both during treatment (p=0.0248) and washout phase (p=0.0036). NASPM in combination with GYKI53655 produced a strong downward trend, which became significant during washout phase (p = 0.0228). Data are shown as scatter plots of individual recordings; superimposed box plots indicate the median (centre line), interquartile range (box edges), and standard deviation (whiskers). <u>Statistics:</u> Kruskal-Wallis with Dunn’s multiple comparisons. Normality assessed via Shapiro-Wilk test. Sample sizes: 0Mg+GBZ+CGP (n=40 slices from 16 patients); +UBP302 (n=11 slices from 7 patients); +GYKI53655 (n=11 slices from 7 patients); +GYKI52466 (n=19 slices from 7 patients); +UBP302+GYKI53655+NBQX (n=22 slices from 10 patients).

The observation that broad AMPAR inhibitors known to also inhibit CP-AMPAR (GYKI52466, GYKI53455 and NBQX, respectively) could reduce, but never abolish, SLE incidence, while the putative specific CP-AMPAR inhibitor NASPM led instead to a complete block of SLE activity, prompted us to ask if other glutamate receptor subtypes could be involved. We first investigated NMDA receptors, another family of glutamatergic receptors permeable to calcium. Similar to the effect of NASPM (0Mg+GBZ+CGP + NASPM: p=0.0030, 6 slices from 6 patients, **Figure 2, D**), application of the NMDA receptor inhibitor APV completely blocked SLE activity induced by 0Mg+GBZ+CGP (p=0.0418, 6 slices from 6 patients, **Figure 5, B**), suggesting that NMDA receptors are necessary for human cortical SLE induced by this method (**Figure 5, A-C**). Due to their strong effect, NASPM and APV were added, differently from the other glutamatergic receptor blockers in study, after SLE were induced. Using NASPM or APV before induction would have hindered to differentiate between its potential inhibitory effect on SLE and to a failure in inducing SLE in the used slice as reported above.

**Figure 5.**
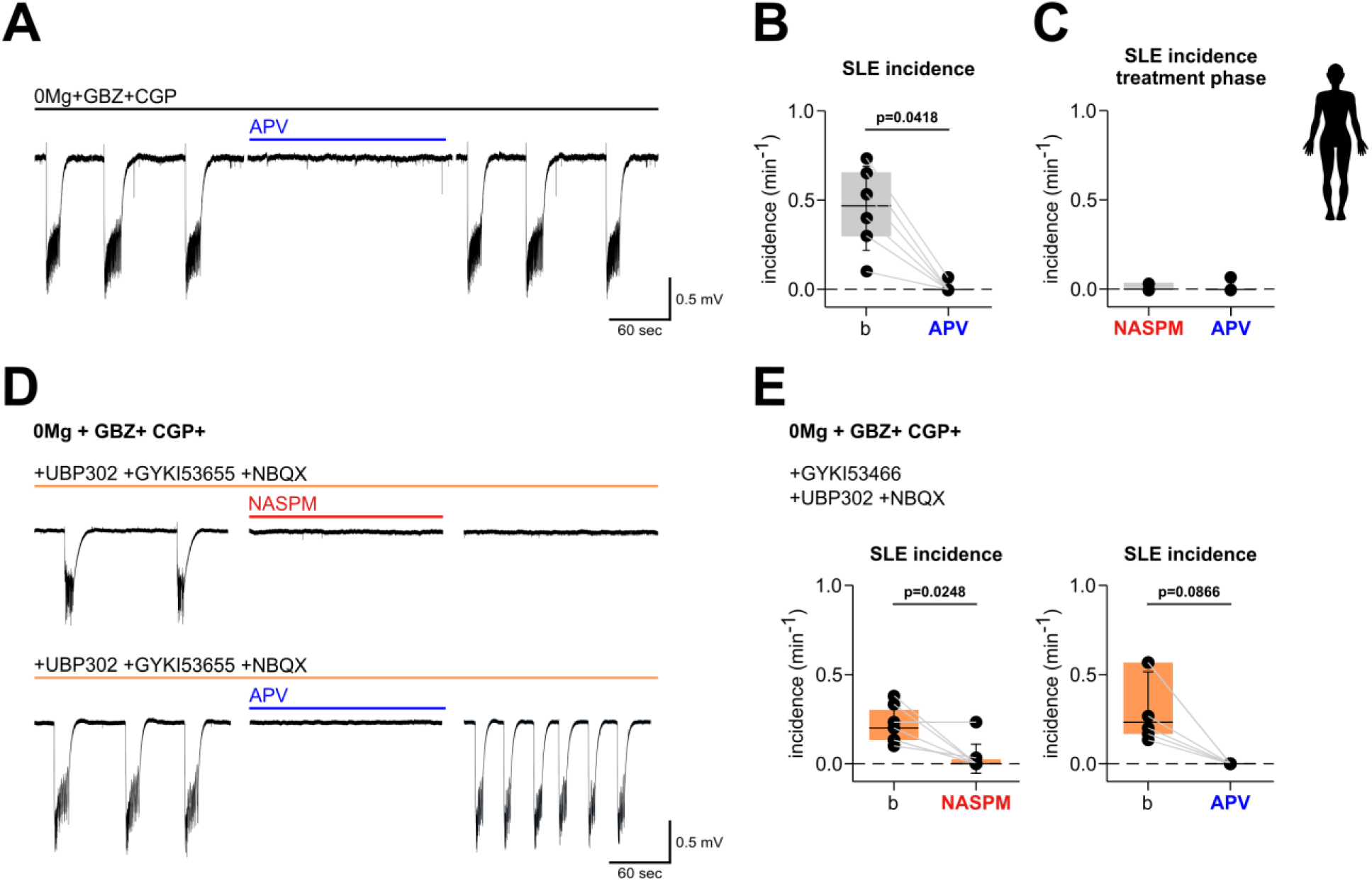
AMPAR/KAR-independent SLEs are blocked by NASPM and APV. **A.** Representative portion of 0Mg+GBZ+CGP baseline, intervention of 50 µM APV and washout. Slow drift removed for clarity. **B.** 50 µM APV application during treatment phase inhibits SLE incidence (p= 0.0418). Data are shown as scatter plots of individual recordings; superimposed box plots indicating the median (centre line), interquartile range (box edges), and standard deviation (whiskers). <u>Statistics</u>: Friedman test with Dunn’s multiple comparisons; normality assessed via Shapiro-Wilk test. **C.** intervention with 100 µM NASPM or 50 µM APV fully inhibits SLE. **D.** SLE present after full combined AMPAR/KAR block are inhibited by APV or NASPM. Representative portion of baseline (0Mg+GBZ+CGP+ 50 µM GYKI53655 + 12.5µM UBP302 + 10 µM NBQX) and intervention of 100 µM NASPM or 50 µM APV show a SLE inhibition upon intervention. SLE fail to restore during analysed time after 100 µM NASPM intervention but are restored after 50 µM APV intervention during washout phase. Slow drift was removed for clarity. **E.** Application of either 100 µM NASPM or 50 µM APV inhibits SLEs induced by 0Mg+GBZ+CGP in the presence of the combined block of KARs and AMPARs (UBP302, GYKI53655 and NBQX). SLE incidence was reduced in a similar manner by 100 µM NASPM (p = 0.0248, n=8 slices from 7 patients) and 50 µM APV (p = 0.0868 n=6 slices from 6 patients). Data are shown as incidence during baseline (b) versus incidence during treatment (NASPM or APV) phase. Data are shown as scatter plots of individual recordings; superimposed box plots indicating the median (centre line), interquartile range (box edges), and standard deviation (whiskers). <u>Statistics</u>: Friedman test with Dunn’s multiple comparisons; normality assessed via Shapiro-Wilk test.

### NASPM blocks seizure-like activity otherwise insensitive to AMPA or kainate receptor inhibitors

Next, we investigated whether the strong inhibitory effects of NASPM and APV on SLE activity were occluded by prior inhibition of AMPA and/or kainate receptors (**Figure 5, E**). In the presence of broad AMPA receptor and kainate receptor inhibition (GYKI53655+UBP302+NBQX), human cortical slices continued to display robust SLE, albeit the incidence was reduced when compared to control (**Figure 4, A**). Subsequent application of 100 µM NASPM or 50 µM APV completely abolished these SLE, strongly suggesting that NASPM may be blocking SLE through inhibition of NMDARs (**Figure 5, E**).

### NASPM inhibits NMDAR mediated currents in human pyramidal neurons

To investigate potential effects of NASPM on NMDA receptors directly, we recorded NMDAR- mediated postsynaptic currents (NMDAR-EPSC) in human brain tissue. We performed single cell patch-clamp recordings in human layer II-III pyramidal neurons from *ex vivo* acute human brain slices. NMDAR-EPSCs were isolated pharmacologically by using a 0Mg+GBZ+CGP+ GYKI53655+UBP302+NBQX induction method (see **Methods**), evoked by extracellular stimulation placed in layer IV/V and recorded at the holding potential of +40 mV to ensure NMDAR activation. During baseline, extracellular stimulation evoked stable EPSC with a slow rise time, suggestive of NMDAR mediated currents. These EPSC showed variable amplitude (442.1 ± 412.8 pA, 22 cells from 11 patients) and charge transfer (area under the curve, 1.15*10^7^ ± 1.3*10^7^, pA*sec, 22 cells from 11 patients). Cell-to-cell variability was attributed to variations of the distance between soma and stimulation electrode and to the network viability. For this reason, data are shown as normalised values (for cell inclusion criteria, see **Methods**). Consistent with our hypothesis, application of 100 µM NASPM strongly reduced amplitude and charge transfer of NMDAR-EPSC (charge transfer reduction: 86.2 % ± 17.8 %; amplitude reduction: 61.7%±33.3%, p= 0.0536; duration reduction: 75.1% ± 23.2%, p=0.0019; time to peak reduction: 75.7%±25.8%, p = 0.0079, 9 cells from 8 patients, **Figure 6, C**). Similarly, currents were fully blocked by 50 µM APV (AUC reduction: 97.58 % ± 2.46%; amplitude reduction: 94.7% ±1.64%; duration reduction: 91.73% ± 4.48%, p = 0.0079, 7 cells from 6 patients; **Figure 6, D**), demonstrating that they were mediated by NMDAR. In control experiments (milli-Q water as vehicle) only the duration was affected slightly (average duration decrease of 24.4% ± 15.2%, 6 cells from 5 patients) while other properties analysed did not vary significantly (**Figure 6)**

**Figure 6.**
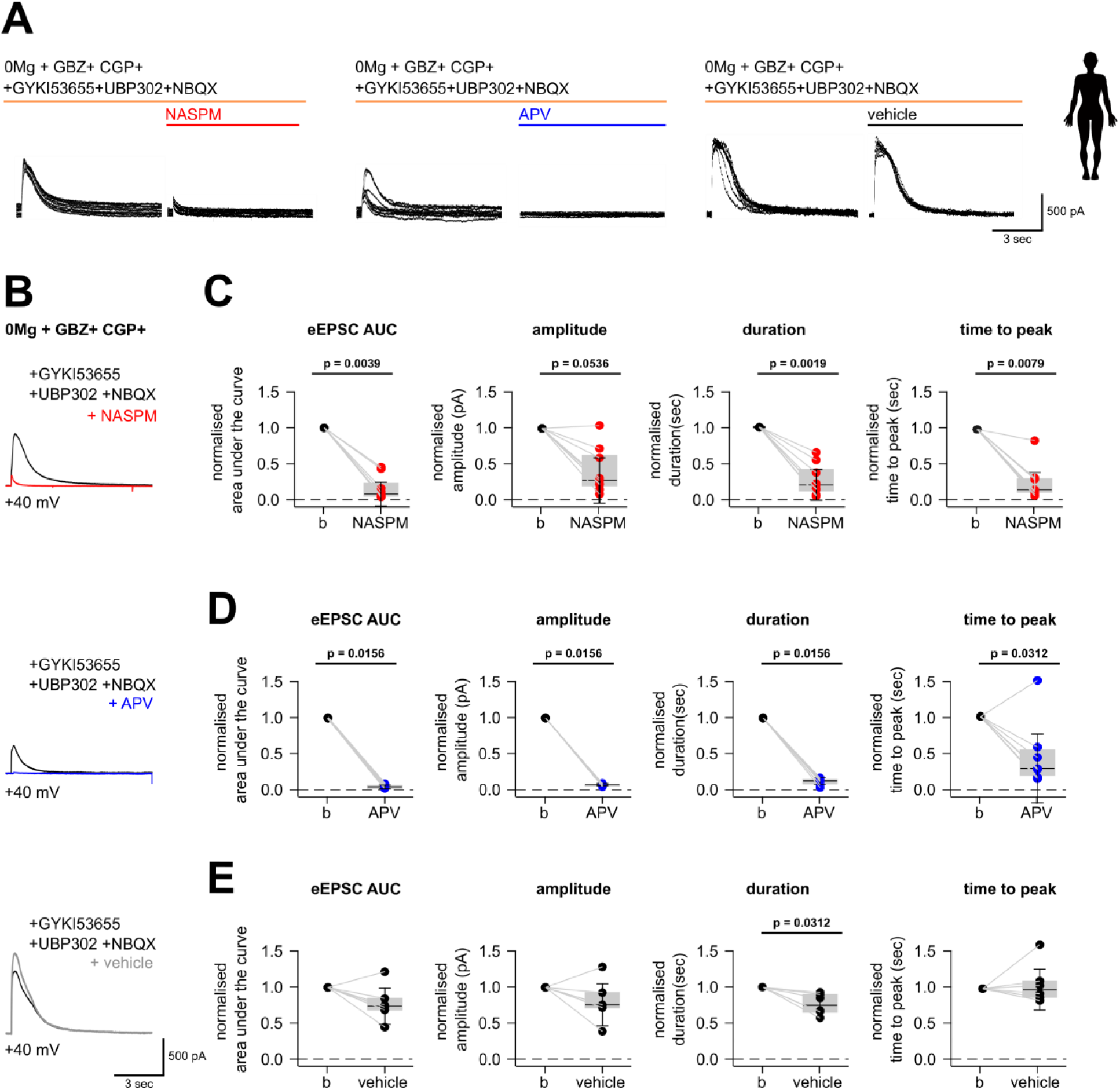
NASPM decreases charge transfer on NMDAR-eEPSCs. **A.** Representative traces of the first 10 sweeps for each condition recorded at +40 mV. **B.** Representative eEPSCs are shown as overlapping traces of 0Mg+GBZ+CGP+GYKI53655+UBP302+NBQX baseline (black) and interventions of 100 µM NASPM (red), 50 µM APV (blue), or 250 µL milli-Q water (vehicle, grey). **C.** NASPM intervention in the 0Mg+GBZ+CGP+GYKI53655+UBP302+NBQX induction model significantly reduced charge transfer (quantified as area under the curve, AUC, p = 0.0039), amplitude (p = 0.0536), duration (p = 0.0019), and time to peak of the amplitude (p = 0.0079, 5 cells from 5 patients). **D.** Similarly, APV intervention significantly decreased eEPSC charge transfer (p = 0.0156), amplitude (p = 0.0156), duration (p = 0.0156) and time to peak (p = 0.0312; 5 cells from 5 patients). Data are shown as scatter plots of normalized values, with superimposed box plots indicating the median (centre line), interquartile range (box edges), and standard deviation (whiskers). <u>Statistics:</u> Mann-Whitney test; normality assessed via Shapiro-Wilk test. **E.** Vehicle intervention mildly altered eEPSC duration (p=0.0312, 6 cells from 6 patients), without significantly altering other eEPSCs analysed properties (p > 0.1, 6 cells from 6 patients).

### NASPM inhibits NMDAR mediated currents in heterologous expression

Our findings in human slices suggested the involvement of NMDA receptors in seizure onset and, surprisingly, their direct inhibition by NASPM under low magnesium conditions. Consequently, we investigated the effect of NASPM directly on heterologously expressed GluN1/GluN2A receptors in whole-cell patch clamp. To ensure reliable solution exchange, we lifted single ND7/23 cells and used a fast perfusion system to apply a magnesium-free extracellular solution containing NMDA/Glycine alone or in combination with NASPM.

Application of NASPM resulted in concentration- and voltage-dependent inhibition of NMDA-evoked currents (**Figure 7, A**). The most pronounced inhibition was observed at –60 mV, yielding normalized peak currents of 7 ± 4%, 16 ± 8%, and 24 ± 5% for 87 µM, 26.1 µM, and 8.7 µM NASPM, respectively. Inhibition at –30 mV was less pronounced, but the peak response was still reduced to 47 ± 14% of control at 26.1 µM NASPM. At +30 mV, NASPM exerted the weakest inhibitory effect (**Figure, 7B**). Notably, the decay of the inhibited responses in continuous NMDA was faster in NASPM, at all voltages in a concentration-dependent manner, in good agreement with the faster residual eEPSCs measured in human slices in the presence of NASPM (**Figure 6**).

**Figure 7.**
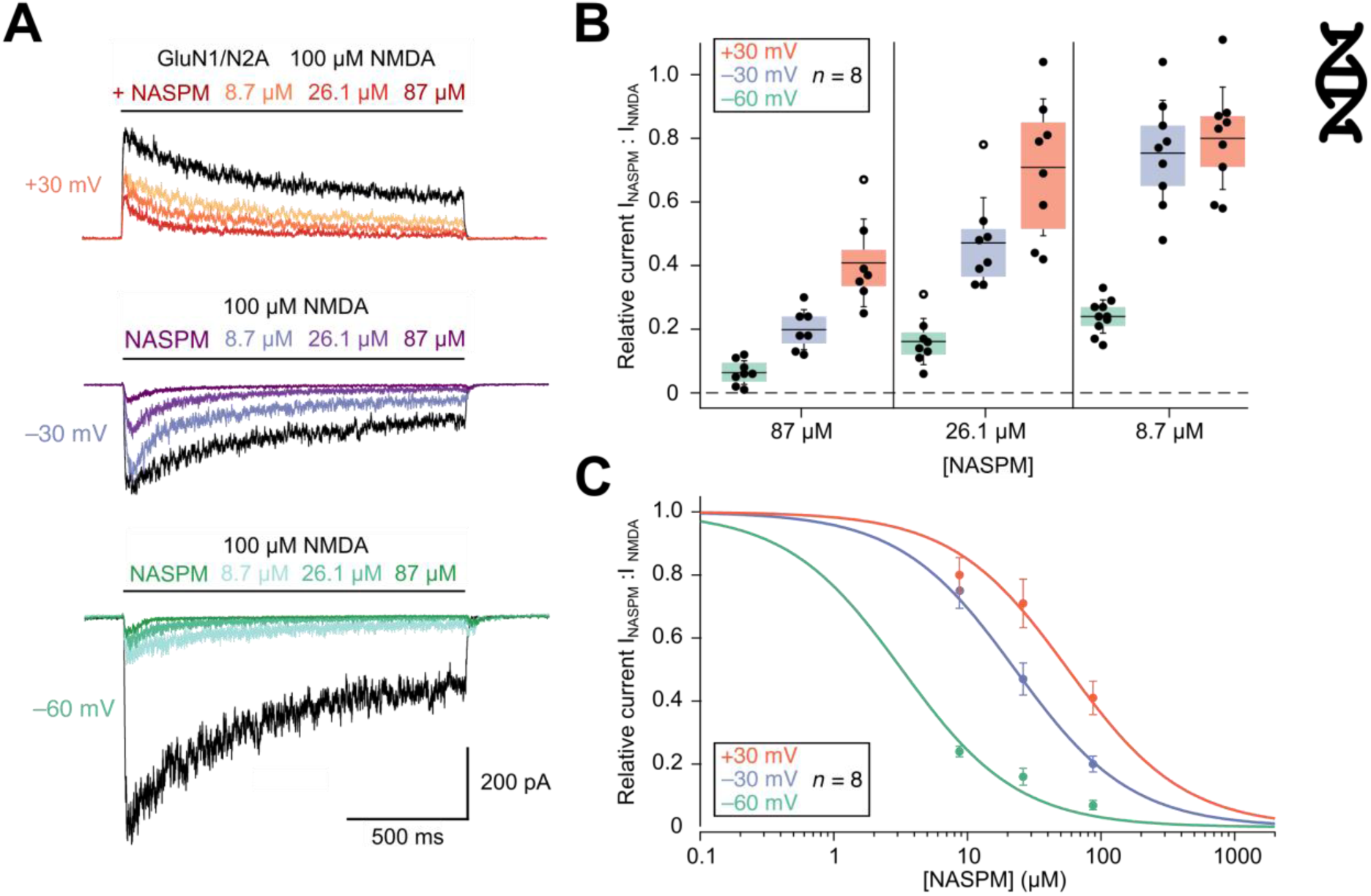
Recombinant GluN1/GluN2A receptors are blocked by NASPM in a concentration- and voltage-dependent manner. **A.** Currents recorded from ND7/23 cells expressing GluN1/GluN2A receptors, at –60 mV, –30 mV and +30 mV, respectively, upon application of NMDA (100 µM, black traces) or NMDA + NASPM (8.7 μM, 26.1 μM, 87 μM, color gradient traces) in a Mg^2+^-free condition. **B.** Normalized GluN1/GluN2A receptor peak currents, categorized by membrane potential and concentration, with upper and lower quartiles, mean and standard deviation. **C.** Estimated concentration dependencies of GluN1/GluN2A block by NASPM. Average normalized GluN1/GluN2A peak currents were fitted with a rectangular hyperbolic function, assuming direct occlusion of the ion channel pore by a single NASPM molecule. The IC_50_ values from the fits were 3.29 ± 0.54 µM, 23.6 ± 1.1 µM, and 56.1 ± 7.4 µM at –60 mV, –30 mV, and +30 mV.

To quantify the potency of NASPM antagonism, normalized peak currents were fitted with a rectangular hyperbolic function assuming direct channel pore occlusion (**Figure, 7C**). The *IC*₅₀ values derived from these fits were 3.29 ± 0.54 µM, 23.6 ± 1.1 µM, and 56.1 ± 7.4 µM at –60 mV, – 30 mV, and +30 mV, respectively, confirming the strong voltage dependence of NASPM-mediated inhibition, and relatively high potency at typical neuronal resting potentials. Even cells depolarized beyond spiking threshold (–30 mV), such as might be found during seizures, were half-blocked by 24 µM NASPM.

### Effects of NASPM on seizure-like activity in mouse tissue

In human cortical tissue, NASPM inhibited SLE activity independently of the induction method, suggesting an essential role of NMDA receptors in human *ex vivo* ictogenesis. To determine whether this effect is species-specific to humans, we investigated the effect of 100 µM NASPM on SLE activity in murine cortical brain slices. Due to differences between human and rodent brain tissue (see **Methods**), SLE in rodent tissue were induced by simpler conditions than in human tissue, either (a) by inhibition of GABA_A_ and GABA_B_ receptors (GBZ+CGP); (b) by disinhibition of NMDA receptors via omission of Mg^2+^ ions in the bath (0Mg) or (c) by pharmacological block of potassium channels (4-AP). When SLE were induced by depletion of Mg ions, NASPM fully and rapidly blocked SLE activity (p=0.0030, **Figure 8, B**), suggesting that it also inhibits NMDA receptors in murine brain. NASPM also inhibited SLE induced by 4AP, although with slower temporal dynamics when compared SLE induced by 0Mg and to SLE in human tissue. Short-term NASPM incubation (< 45 min) partially blocked murine SLE induced by 4AP, whereas prolonged incubation with NASPM (90 min) fully suppressed SLEs (p=0.0145; 5 slices from 5 animals, **Figure 8, A-B**). When SLE were induced by blocking GABAergic transmission (GBZ+CGP), the resulting events were short in duration and more frequent, resembling bursting activity rather than SLE. NASPM had no effect on the incidence or properties of these events (**Figure 8, A-B**). Taken together, the inhibitory effect of 100 µM NASPM on SLE was observed in murine tissue, as in human cortical slices.

**Figure 8.**
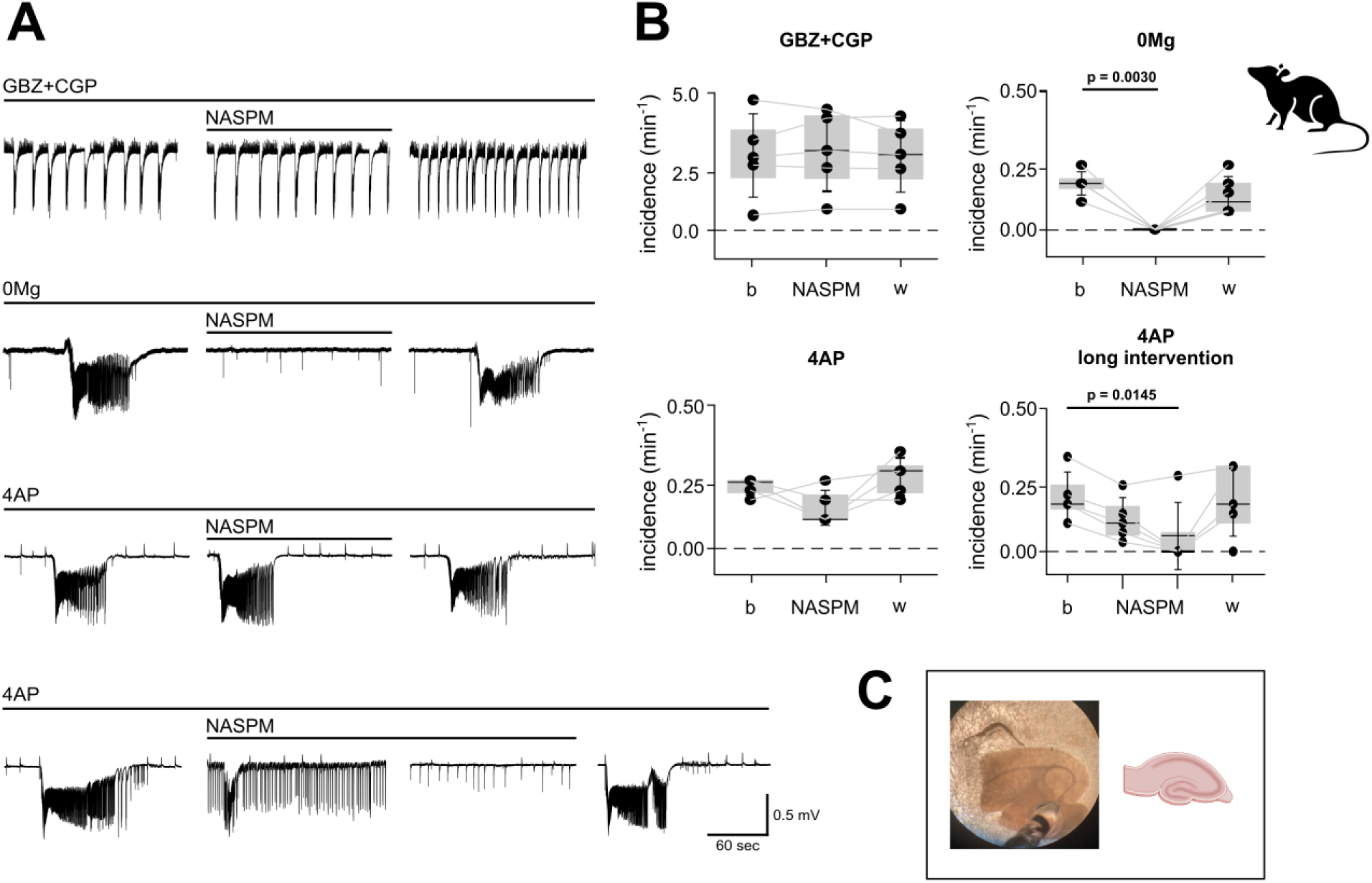
NASPM effects on bursting activity and SLEs in ex vivo mouse brain tissue. **A.** In mouse brain slices, NASPM intervention on SLE incidence depended on the induction method and exposure time. Representative portions of baseline, NASPM intervention and washout phases are shown for each method of induction. Induction by GBZ+CGP induced high frequency epileptic activity rather than SLEs. Slow baseline drift of traces was removed for better visualisation. **B.** Quantification of NASPM intervention on SLE incidence across various induction methods. High frequency epileptic activity induced by GBZ+CGP induction method is not affected by intervention of 100 µM NASPM (left upper panel, 5 slices from 5 animals); in contrast, NASPM significantly blocks SLE incidence when a 0Mg induction method is used to induce SLE (p= 0.0030, 6 slices from 6 animals, right-upper panel). 100 µM NASPM does not significantly reduce SLE incidence upon 4AP induction (left lower panel, 5 slices from 5 animals), but did induce a significant decrease when a longer intervention period is maintained (right lower panel, p= 0.0145. 5 slices from 5 animals). Data are shown as scatter plots of individual recordings; superimposed box plots indicate the median (centre line), interquartile range (box edges), and standard deviation (whiskers). <u>Statistics</u>: Friedman test with Dunn’s multiple comparison test. Normality was calculated with Shapiro-Wilk test. **C.** Micrograph of a representative 450µm thick hippocampal slice used in these experiments.

## DISCUSSION

In this study, we examined the role of ionotropic glutamatergic receptor subtypes in seizure-like activity induced in human cortical tissue *ex vivo*. Our key findings are as follows: (1) using 0Mg+GBZ+CGP was the most effective method for inducing SLE in *ex vivo* human cortex, although the incidence and biophysical properties of SLE were similar between induction methods tested. (2) The CP-AMPAR blocker NASPM inhibited SLE in human tissue regardless of the induction method, while IEM-1460, another CP-AMPAR inhibitor, did not suppress SLE. (3) Several AMPA and kainate receptor antagonists, either alone or in combination, reduced the incidence of SLE but did not abolish it, whereas the intervention of NMDA receptor antagonist APV, fully blocked seizure like activity. (4) SLE resistant to AMPA and/or kainate receptor inhibition was completely suppressed by NASPM or APV, suggesting a shared mechanism of action. This finding was further supported by experiments showing that (5) NASPM inhibited NMDAR-mediated EPSC in human cortical neurons and (6) heterologously-expressed NMDAR currents. (7) NASPM also suppressed SLE in *ex vivo* rodent cortical slices, demonstrating that its effect is conserved. Our results support the essential role of NMDA receptors in ictogenesis in human tissue and, most importantly, our findings challenge the view of NASPM as a specific CP-AMPAR inhibitor.

### Ionotropic glutamatergic receptors in ictogenesis and epileptogenesis

Our results demonstrate that AMPAR and KAR contribute independently to SLE but confirm the essential role of NMDAR in SLE generation while finding no clear evidence for the specific involvement of CP-AMPAR in human SLE *ex vivo*. CP-AMPAR have been suggested to contribute to neuronal dysfunction and excitotoxicity in epilepsy due to the greater Ca²⁺ influx through unedited GluA2 subunits. This conceptual frame has been proposed to link CP-AMPAR expression to ictogenesis (Guo et al., 2021; Higuchi et al., 2000; Brusa et al., 1995). Targeting CP-AMPAR has therefore been explored as a potential therapeutic approach with promising results in animal models (Zinchenko et al., 2024; Noh et al., 2005; Kanai et al., 1992). In murine models, CP-AMPAR are predominantly expressed in fast-spiking inhibitory interneurons (Lalanne et al., 2016; Lalanne et al., 2018), suggesting that their inhibition would reduce interneuron activity and shift the excitation-inhibition balance toward excitation, unless GABA receptors are excitatory due to altered intracellular Cl⁻ (McMoneagle et al., 2024; van Hugte et al., 2023; Miles et al., 2021; Huberfeld et al., 2007). Following this logic, SLE in the presence of GABAergic inhibition would not expect to be affected by CP-AMPAR suppression, unless CP-AMPAR are expressed in excitatory neurons. Although some data suggest that CP-AMPAR could be expressed in hippocampal principal cells, particularly during plasticity, there is little reason to think that specific inhibition of CP-AMPAR in excitatory neurons would be decisive for SLE (Kochlamazashvili et al., 2022; Sanderson et al., 2016). In our hands, the two polyamine drugs taken to be selective CP-AMPAR antagonists, IEM-1460 and NASPM, exhibited divergent effects. IEM-1460 did not affect SLE incidence in the presence of GABAergic inhibition, as expected. In contrast, NASPM fully blocked SLE, however, this effect was almost certainly mediated by non-specific inhibition of NMDA receptors as discussed below, and not dependent on CP-AMPARs.

Similar to human tissue, ictogenesis in murine tissue was also dependent on NMDAR activation, suggesting that mechanisms underlying cortical ictogenesis *in vitro* show some conservation across species. However, it is important to note that our experiments were conducted exclusively in "seizure-conditioned" human cortical tissue *ex vivo*, whereas the murine tissue was from healthy mice, and so these findings cannot be easily generalized to epilepsy mechanisms in the ictogenic zone (e.g., hippocampus in temporal lobe epilepsy) or in intact *in vivo* networks. NMDAR have been shown to play a critical role in seizure onset (Gjerulfsen et al., 2024; Hanson et al., 2024; Sadeghi et al., 2021; Hanada, 2020; Husari et al., 2019; Cohen et al., 2002; Rogawski et al., 1992), and NMDAR antagonists have been explored as therapeutic targets in preclinical trials (Ghasemi and Schachter, 2011) or as part of approved anti-seizure treatments (Harty et al., 2000; Kleckner et al., 1999). However, targeting NMDAR presents challenges, as their inhibition can lead to undesirable effects as seen in patients with NMDAR autoantibodies (Husari et al., 2019; Miya et al., 2014). Overall, we attribute the effects of NASPM primarily to its inhibition of NMDAR, and since IEM-1460 showed no clear effects, we currently see no evidence for a primary involvement of CP-AMPAR in human ictogenesis.

### Species Differences in SLE Induction and Temporal Dynamics

SLE induction thresholds and temporal dynamics of SLE differed between human and murine tissue. In particular, in human tissue SLE threshold was markedly higher and consistent with previous observations (Peng et al., 2024; Kraus et al., 2020; Schwarz et al., 2017; Eugène et al., 2014; Wahab et al., 2010; Gabriel et al., 2004), requiring the combination of two or more pharmacological treatments. These differences may be attributed to cross-species variations in cellular density, dendritic arborization, and intrinsic excitability (Chartrand et al., 2023; Lee et al., 2023; Wilbers et al., 2023). Additionally, the temporal dynamics of NASPM effects varied across induction methods in murine tissue, likely due to differences in the NMDAR pore state; while in 0Mg, with NMDAR pore persistently open NASPM exerted rapid effects, in 4AP, NASPM effect was delayed. These differences represent a limitation and should be considered when interpreting data from *in vitro* models across species or using different induction paradigms.

### NASPM effect on NMDAR

Our data together with a literature search imply that the inhibitory effect of NASPM on NMDAR has been underappreciated for decades. NASPM is a synthetic analogue of Joro spider toxin (JSTX), first characterized in 1989 (Asami et al., 1989); JSTX and its analogues, including NASPM, were shown to block CP-AMPAR both in heterologous expression (Washburn and Dingledine, 1994; Blaschke et al., 1993; Herlitze et al., 1993) and in rodent CNS (Koike et al., 1997; Gu et al., 1996; Iino et al., 1996; Meucci et al., 1996). Among JSTX analogues, NASPM has been considered the most promising for research and putative clinical application due to its high affinity for CP-AMPAR and reversible mechanism of action (Twomey et al., 2018; Asami et al., 1989). Interestingly, CP-AMPAR are not the exclusive targets of polyamines. The presence of an allosteric binding site for the endogenous polyamine spermine on NMDAR was firstly hypothesised in 1992 (Lerma et al., 1992). Both endogenous and synthetic polyamines have been known for long time to exert effects on NMDARs (Mony et al., 2011; Williams et al., 1997; Munir et al., 1993; Lerma et al., 1992; Brackley et al., 1990; Reynolds et al., 1990), but that these effects could implicate NASPM as an NMDA blocker appears to have drifted from view. NASPM was described to exhibit low affinity for NMDAR (Isaac et al., 2007; Washburn and Dingledine, 1996), and similar compounds, such as ArgTX-636 (Usherwood et al., 1991; Priestley et al., 1989) and JSTX-3 (Salamoni et al., 2005; Mueller et al., 1991) have been shown to inhibit NMDAR activity.

While previous studies demonstrated NASPM inhibitory effect on CP-AMPAR, often potential effects on NMDARs were not investigated. Nevertheless, many studies have since referred to NASPM as a specific CP-AMPAR inhibitor. In these studies, cells were often clamped at –65±5 mV, a potential that does not exclude NMDAR activation due to partial NMDA activity at this potential and space-clamp differences between soma and dendrites. Additionally, the extracellular Mg²⁺ concentrations used (1-2 mM) in most studies were insufficient to fully block NMDAR, leaving the receptors partially open and susceptible to NASPM action even at –60 mV (Wang et al., 2004). Recently, it was recognized that extracellular magnesium levels are perhaps lower than previously thought, 0.7 mM, and do not fully inhibit NMDA receptors at rest (Chiu et al., 2022). However, for studies that were not performed in voltage clamp on single cells, membrane potentials could also be depolarised leading to a much stronger inhibition by NASPM (**Figure 7**). Particularly, this concern applies whenever NASPM was used *in vivo,* and many studies did not perform controls to exclude NMDAR inhibition when using NASPM. We believe that the widespread use of NASPM as a specific CP-AMPAR inhibitor stems from a misunderstanding of its initially described effects on AMPAR, with good evidence that **NASPM is specific for CP-AMPARs within the AMPAR family** (Iino et al., 1996; Blaschke et al., 1993). However, this specificity has been misinterpreted, leading to the misconception that NASPM does not affect other glutamate receptors. In contrast to its widespread use as a specific CP-AMPAR inhibitor, our study provides clear evidence that NASPM also acts as an inhibitor of NMDA receptors in both human and murine tissue.

## Conclusion

Overall, our study highlights a long-misunderstood lack of specifity of NASPM, and prompt reassessment of NASPM’s dual action on CP-AMPA receptors and NMDA receptors. The results from human tissue indicate promise for polyamine-like molecules for block of SLE. Further work will be needed to clarify to the pharmacology of glutamate receptors, and particularly the specificity of different synthetic polyamines like IEM-1460 and NASPM for glutamatergic receptor subtypes.

## Data availability

Raw data generated during this study are available upon request to the corresponding authors.

## Ethics statements

Experiments were conducted in accordance with the German Animal Welfare Act and the European Directive 2010/63/EU for animal experiments, and with approval of the Institutional Animal Welfare Officer and the responsible local authorities (Landesamt für Gesundheit und Soziales Berlin, T0336/12).

All patients provided informed consent for tissue donation. Human derived material was treated in accordance with and the study was positively assessed by the local ethical committee (Berlin: vote no. EA2/111/14, ethical commission of the Charité, Bielefeld: vote no.2020-517-f-S, ethical commission of the medical chamber Westfalen Lippe, Hamburg: vote no. 2023-200674-BO-bet, ethical commission of the medical chamber Hamburg).

## Author contributions

PF, AJRP, JRPG and MH were involved in study design and conception. RX, JO, U-WT, TK, MS and TS informed patients to get consent and contributed to surgical resection of the tissue material used in this study. JA, AJRP and SB designed, performed and analysed the electrophysiological experiments in the heterologous expression system. PF, HA and JRPG coordinated cooperations and obtained ethical approvals. AP performed electrophysiological recordings in mouse and human *ex vivo* model. LM and FA contributed to execution of electrophysiological experiments in human ex vivo slices. AP analysed the data and performed the statistical analysis. AP wrote the first draft of the manuscript and performed the data visualisation. PF revised the manuscript. All authors read and approved the final version of the manuscript.

## Acknowledgments

This work is part of the doctoral thesis of AP. We would like to thank Mandy Marbler-Pötter and Andrea Wilke for their excellent technical assistance. We thank Laetitia Mony (ENS, Paris) for advice and for providing pilot experiments on NMDA sensitivity to NASPM. The resources for this project were provided by institutional funding of the Institute of Neurophysiology and of the Department of Experimental Neurology, both part of Charité – Universitätsmedizin Berlin, corporate member of Freie Universität Berlin and Humboldt Universität zu Berlin (JG, MH). This research was funded by the Deutsche Forschungsgemeinschaft under Germany’s Excellence Strategy (EXC-2049-390688087-NeuroCure) to A.J.R.P.

## Abbreviations

4AP: 4 -Aminopyridine
aCSF: artificial cerebrospinal fluid
AMPARs: Alpha-Amino-3-hydroxy-5-methylisoxazole-4-propionate receptors
APV: D-(-)-2-Amino-5-phosphonopentanoic acid
b: baseline
BIC: bicuculline
CP-AMPAR: Ca^2+^ - permeable AMPARs
eEPSCs: evoked excitatory postsynaptic currents
EPSCs: excitatory postsynaptic currents
KARs: kainate receptors
GABA: Gamma-aminobutyric acid
GBZ: gabazine
JSTX: Joro spider toxin
LFP: local field potential
MM: Materials and Methods
NASPM: naphthyl-acetyl spermine trihydrochloride
NMDARs: N-methyl-D-aspartate receptors
Ra: access resistance
Rm: membrane resistance
Rs: series resistance
w: washout

---

**Supplementary Figure 1.**
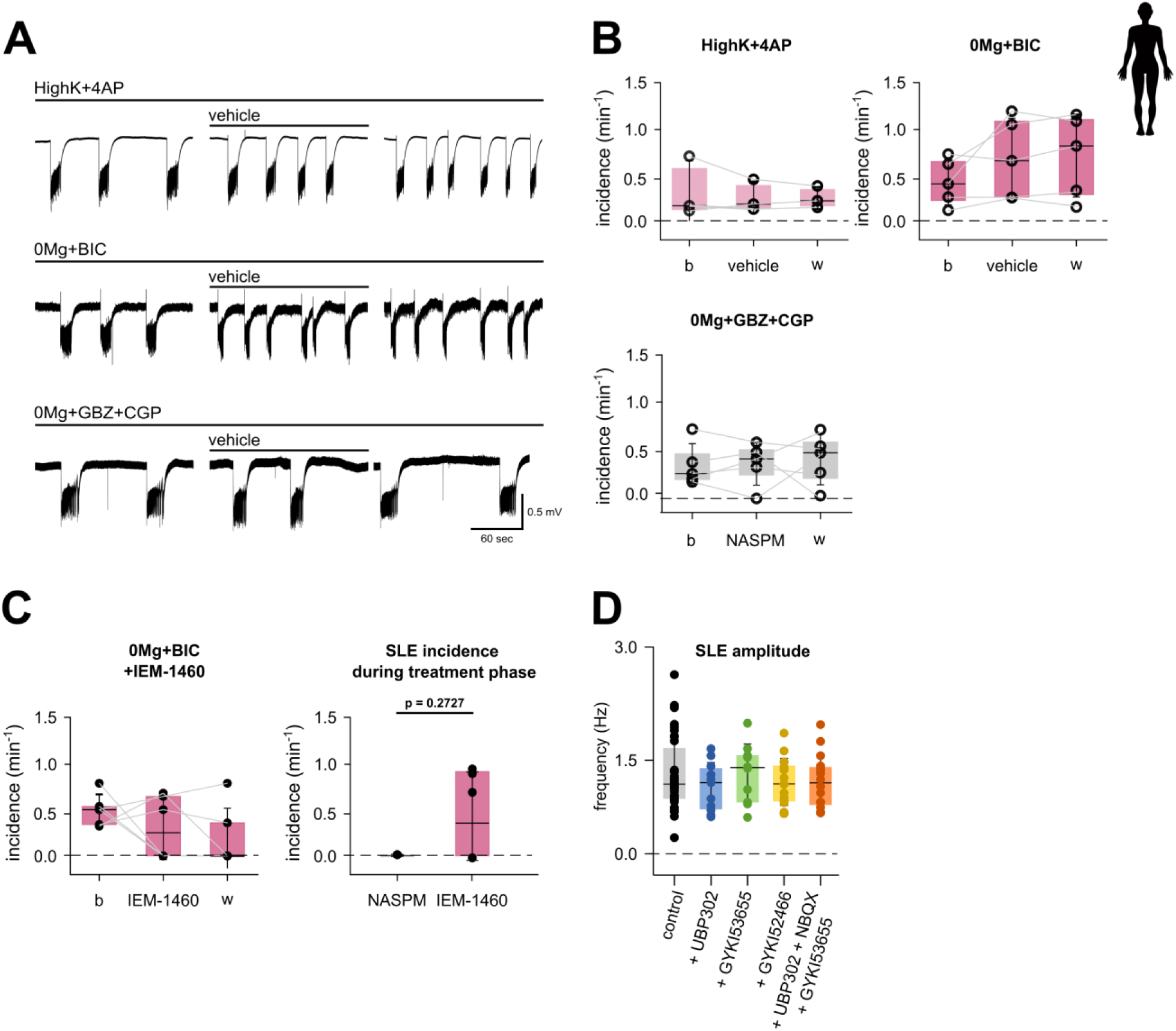
Vehicle or IEM-1460 intervention does not affect SLEs incidence. SLE amplitude and duration is not affected by glutamatergic blockers UBP302, GYKI53655, GYKI5245, NBQX. **A.** Vehicle intervention does not significantly reduce SLEs incidence during intervention phase. For each induction method a representative portion of baseline, vehicle intervention and washout phase are shown, slow drift of the traces were removed for better visualization **B.** SLE incidence remains stable over time during the intervention of 300 µL Milli-Q water as vehicle in all induction methods studied (High+4AP, 5 slices from 3 patients; 0Mg+BIC, 7 slices from 5 patients; 0Mg+GBZ+CGP, 6 slices from 5 patients). <u>Statistics</u>: Friedman test with Dunn’s multiple comparisons; normality was assessed via Shapiro-Wilk test. **C.** Similarly, IEM-1460 intervention does not affect significantly SLEs incidence in a 0Mg+BIC induction method (left panel). <u>Statistics</u>: Friedman test with Dunn’s multiple comparisons; normality assessed via Shapiro-Wilk test. A significant difference in observed in SLE incidence during intervention of 100 µM NASPM (9 slices from 5 patients) or 100µM IEM-1460 (9 slices from 6 patients, p = 0.2727, right panel). <u>Statistics</u>: Mann-Whitney test; normality was assessed via Shapiro-Wilk test. All data are shown as scatter plots of individual recordings; superimposed box plots indicate the median (centre line), interquartile range (box edges), and standard deviation (whiskers). **D.** In the induction method 0Mg+GBZ+CGP with addition of various glutamatergic UBP302, GYKI53655, GYKI52455, and a combination of UBP302, GYKI53655, and NBQX, SLE amplitude and duration doesńt vary significantly across the methods. 0Mg+GBZ+CGP+ NASPM and +APV as intervention compounds are shown for completeness. <u>Statistics:</u> Kruskal-Wallis with Dunn’s multiple comparisons. Normality assessed via Shapiro-Wilk test. Sample sizes: 0Mg+GBZ+CGP (n=40 slices from 16 patients); +UBP302 (n=11 slices from 7 patients); +GYKI53655 (n=11 slices from 7 patients); +GYKI52466 (n=19 slices from 7 patients); +UBP302+GYKI53655+NBQX (n=22 slices from 10 patients).

## REFERENCES

1. Agarwal, R., & Smith, J. C. (2023). Speed vs accuracy: effect on ligand pose accuracy of varying box size and exhaustiveness in AutoDock vina. Molecular Informatics, 42(2), 2200188.

2. Asami, T., Kagechika, H., Hashimoto, Y., Shudo, K., Miwa, A., Kawai, N., & Nakajima, T. (1989). Acylpolyamines mimic the action of joro spider toxin (JSTX) on crustacean muscle glutamate receptors. Biomedical research, 10(3), 185–189.

3. Barria, A., & Malinow, R. (2002). Subunit-specific NMDA receptor trafficking to synapses. Neuron, 35(2), 345–353.

4. Blaschke, M., Keller, B. U., Rivosecchi, R., Hollmann, M., Heinemann, S., & Konnerth, A. (1993). A single amino acid determines the subunit-specific spider toxin block of alpha-amino-3-hydroxy-5-methylisoxazole-4-propionate/kainate receptor channels. Proceedings of the National Academy of Sciences, 90(14), 6528–6532.

5. Bonansco, C., & Fuenzalida, M. (2016). Plasticity of hippocampal excitatory-inhibitory balance: Missing the synaptic control in the epileptic brain. Neural plasticity, 2016(1), 8607038.

6. Boulter, J., Hollmann, M., O’Shea-Greenfield, A., Hartley, M., Deneris, E., Maron, C., & Heinemann, S. (1990). Molecular cloning and functional expression of glutamate receptor subunit genes. Science, 249(4972), 1033–1037.

7. Bowie, D., & Mayer, M. L. (1995). Inward rectification of both AMPA and kainate subtype glutamate receptors generated by polyamine-mediated ion channel block. Neuron, 15(2), 453–462.

8. Brusa, R., Zimmermann, F., Koh, D. S., Feldmeyer, D., Gass, P., Seeburg, P. H., & Sprengel, R. (1995). Early-onset epilepsy and postnatal lethality associated with an editing-deficient GluR-B allele in mice. Science, 270(5242), 1677–1680.

9. Burnashev, N., Monyer, H., Seeburg, P. H., & Sakmann, B. (1992). Divalent ion permeability of AMPA receptor channels is dominated by the edited form of a single subunit. Neuron, 8(1), 189–198.

10. Chartrand, T., Dalley, R., Close, J., Goriounova, N. A., Lee, B. R., Mann, R., … & Lein, E. S. (2023). Morphoelectric and transcriptomic divergence of the layer 1 interneuron repertoire in human versus mouse neocortex. Science, 382(6667), eadf0805.

11. Chiu, D. N., & Carter, B. C. (2022). Synaptic NMDA receptor activity at resting membrane potentials. Frontiers in Cellular Neuroscience, 16, 916626.

12. Cohen, I., Navarro, V., Clemenceau, S., Baulac, M., & Miles, R. (2002). On the origin of interictal activity in human temporal lobe epilepsy in vitro. Science, 298(5597), 1418–1421.

13. Diering, G. H., & Huganir, R. L. (2018). The AMPA receptor code of synaptic plasticity. Neuron, 100(2), 314–329.

14. Eberhardt, J., Santos-Martins, D., Tillack, A. F., & Forli, S. (2021). AutoDock Vina 1.2. 0: New docking methods, expanded force field, and python bindings. Journal of chemical information and modeling, *61*(8), 3891-3898.

15. Eugène, E., Cluzeaud, F., Cifuentes-Diaz, C., Fricker, D., Le Duigou, C., Clemenceau, S., … & Miles, R. (2014). An organotypic brain slice preparation from adult patients with temporal lobe epilepsy. Journal of Neuroscience Methods, 235, 234–244.

16. Forli, S., Huey, R., Pique, M. E., Sanner, M. F., Goodsell, D. S., & Olson, A. J. (2016). Computational protein–ligand docking and virtual drug screening with the AutoDock suite. Nature protocols, 11(5), 905–919.

17. Ghasemi, M., & Schachter, S. C. (2011). The NMDA receptor complex as a therapeutic target in epilepsy: a review. Epilepsy & Behavior, 22(4), 617–640.

18. Gjerulfsen, C. E., Krey, I., Klöckner, C., Rubboli, G., Lemke, J. R., & Møller, R. S. (2024). Spectrum of NMDA Receptor Variants in Neurodevelopmental Disorders and Epilepsy. NMDA Receptors: Methods and Protocols, 1-11.

19. Gabriel, S., Njunting, M., Pomper, J. K., Merschhemke, M., Sanabria, E. R., Eilers, A., … & Lehmann, T. N. (2004). Stimulus and potassium-induced epileptiform activity in the human dentate gyrus from patients with and without hippocampal sclerosis. Journal of Neuroscience, 24(46), 10416–10430.

20. Geiger, J. R., Melcher, T., Koh, D. S., Sakmann, B., Seeburg, P. H., Jonas, P., & Monyer, H. (1995). Relative abundance of subunit mRNAs determines gating and Ca2+ permeability of AMPA receptors in principal neurons and s in rat CNS. Neuron, 15(1), 193–204.

21. Greger, I. H., Watson, J. F., & Cull-Candy, S. G. (2017). Structural and functional architecture of AMPA-type glutamate receptors and their auxiliary proteins. Neuron, 94(4), 713–730.

22. Guo, C., & Ma, Y. Y. (2021). Calcium permeable-AMPA receptors and excitotoxicity in neurological disorders. Frontiers in neural circuits, 15, 711564.

23. Hanada, T. (2020). Ionotropic glutamate receptors in epilepsy: a review focusing on AMPA and NMDA receptors. Biomolecules, 10(3), 464.

24. Hansen, Kasper B., Lonnie P. Wollmuth, Derek Bowie, Hiro Furukawa, Frank S. Menniti, Alexander I. Sobolevsky, Geoffrey T. Swanson et al. "Structure, function, and pharmacology of glutamate receptor ion channels." Pharmacological reviews 73, no. 4 (2021): 1469–1658.

25. Hanson, J. E., Yuan, H., Perszyk, R. E., Banke, T. G., Xing, H., Tsai, M. C., … & Traynelis, S. F. (2024). Therapeutic potential of N-methyl-D-aspartate receptor modulators in psychiatry. Neuropsychopharmacology, 49(1), 51–66.

26. Harty, T. P., & Rogawski, M. A. (2000). Felbamate block of recombinant N-methyl-D-aspartate receptors: selectivity for the NR2B subunit. Epilepsy research, 39(1), 47–55.

27. Herlitze, S., Raditsch, M., Ruppersberg, J. P., Jahn, W., Monyer, H., Schoepfer, R., & Witzemann, V. (1993). Argiotoxin detects molecular differences in AMPA receptor channels. Neuron, 10(6), 1131–1140.

28. Higuchi, M., Single, F. N., Köhler, M., Sommer, B., Sprengel, R., & Seeburg, P. H. (1993). RNA editing of AMPA receptor subunit GluR-B: a base-paired intron-exon structure determines position and efficiency. Cell, 75(7), 1361–1370.

29. Higuchi, M., Maas, S., Single, F. N., Hartner, J., Rozov, A., Burnashev, N., … & Seeburg, P. H. (2000). Point mutation in an AMPA receptor gene rescues lethality in mice deficient in the RNA-editing enzyme ADAR2. Nature, 406(6791), 78–81.

30. Hsiao, M. C., Yu, P. N., Song, D., Liu, C. Y., Heck, C. N., Millett, D., & Berger, T. W. (2015). An in vitro seizure model from human hippocampal slices using multi-electrode arrays. Journal of neuroscience methods, 244, 154–163.

31. Huberfeld, G., Wittner, L., Clemenceau, S., Baulac, M., Kaila, K., Miles, R., & Rivera, C. (2007). Perturbed chloride homeostasis and GABAergic signaling in human temporal lobe epilepsy. Journal of Neuroscience, 27(37), 9866–9873.

32. Husari, K. S., & Dubey, D. (2019). Autoimmune epilepsy. Neurotherapeutics, 16(3), 685–702.

33. Isaac, J. T., Ashby, M. C., & McBain, C. J. (2007). The role of the GluR2 subunit in AMPA receptor function and synaptic plasticity. Neuron, 54(6), 859–871.

34. Iino, M., Koike, M., Isa, T., & Ozawa, S. (1996). Voltage-dependent blockage of Ca (2+)-permeable AMPA receptors by joro spider toxin in cultured rat hippocampal neurones. The Journal of Physiology, 496(2), 431–437.

35. Isaac, J. T., Ashby, M. C., & McBain, C. J. (2007). The role of the GluR2 subunit in AMPA receptor function and synaptic plasticity. Neuron, 54(6), 859–871.

36. Lalanne, T., Oyrer, J., Mancino, A., Gregor, E., Chung, A., Huynh, L., … & Sjöström, P. J. (2016). Synapse-specific expression of calcium-permeable AMPA receptors in neocortical layer 5. The Journal of Physiology, 594(4), 837–861.

37. Lalanne, T., Oyrer, J., Mancino, A., Gregor, E., Chung, A., Huynh, L., … & Sjöström, P. J. (2016). Synapse-specific expression of calcium-permeable AMPA receptors in neocortical layer 5. The Journal of Physiology, 594(4), 837–861.

38. Lee, B. R., Dalley, R., Miller, J. A., Chartrand, T., Close, J., Mann, R., … & Ting, J. T. (2023). Signature morphoelectric properties of diverse GABAergic interneurons in the human neocortex. Science, 382(6667), eadf6484.

39. Noh, K. M., Yokota, H., Mashiko, T., Castillo, P. E., Zukin, R. S., & Bennett, M. V. (2005). Blockade of calcium-permeable AMPA receptors protects hippocampal neurons against global ischemia-induced death. Proceedings of the National Academy of Sciences, 102(34), 12230–12235.

40. Jonas, P., Racca, C., Sakmann, B., Seeburg, P. H., & Monyer, H. (1994). Differences in Ca2+ permeability of AMPA-type glutamate receptor channels in neocortical neurons caused by differential GluR-B subunit expression. Neuron, 12(6), 1281–1289.

41. Kanai, H., Ishida, N., Nakajima, T., & Kato, N. (1992). An analogue of Joro spider toxin selectively suppresses hippocampal epileptic discharges induced by quisqualate. Brain research, 581(1), 161–164.

42. Kawahara, Y., & Kwak, S. (2005). Excitotoxicity and ALS: what is unique about the AMPA receptors expressed on spinal motor neurons?. Amyotrophic Lateral Sclerosis, 6(3), 131–144.

43. Kleckner, et al., 1999;, N. W., Glazewski, J. C., Chen, C. C., & Moscrip, T. D. (1999). Subtype-selective antagonism of N-methyl-D-aspartate receptors by felbamate: insights into the mechanism of action. The Journal of pharmacology and experimental therapeutics, *289*(2), 886-894. i.e. carbamazepine or felbamate

44. Kochlamazashvili, G., Pampaloni, N. P., Sposini, S., Shergill, J. K., Lehmann, M., Pashkova, N., … & Maritzen, T. (2022). Selective endocytosis of Ca 2+-permeable AMPARs by the Alzheimer’s disease risk factor CALM bidirectionally controls synaptic plasticity. Science Advances, 8(21), eabl5032-eabl5032.

45. Koh, D. S., Geiger, J. R., Jonas, P., & Sakmann, B. (1995). Ca (2+)-permeable AMPA and NMDA receptor channels in basket cells of rat hippocampal dentate gyrus. The Journal of physiology, 485(2), 383–402.

46. Koike, M., Iino, M., & Ozawa, S. (1997). Blocking effect of 1-naphthyl acetyl spermine on Ca2+-permeable AMPA receptors in cultured rat hippocampal neurons. Neuroscience research, 29(1), 27–36.

47. Konen, L. M., Wright, A. L., Royle, G. A., Morris, G. P., Lau, B. K., Seow, P. W., … & Vissel, B. (2020). A new mouse line with reduced GluA2 Q/R site RNA editing exhibits loss of dendritic spines, hippocampal CA1-neuron loss, learning and memory impairments and NMDA receptor-independent seizure vulnerability. Molecular brain, 13, 1–19.

48. Kortenbruck, G., Berger, E., Speckmann, E. J., & Musshoff, U. (2001). RNA editing at the Q/R site for the glutamate receptor subunits GLUR2, GLUR5, and GLUR6 in hippocampus and temporal cortex from epileptic patients. Neurobiology of disease, *8*(3), 459-468.

49. Kromann, H., Krikstolaityte, S., Andersen, A. J., Andersen, K., Krogsgaard-Larsen, P., Jaroszewski, J. W., … & Strømgaard, K. (2002). Solid-phase synthesis of polyamine toxin analogues: potent and selective antagonists of Ca2+-permeable AMPA receptors. Journal of medicinal chemistry, 45(26), 5745–5754.

50. Kumar, S. S., Bacci, A., Kharazia, V., & Huguenard, J. R. (2002). A developmental switch of AMPA receptor subunits in neocortical pyramidal neurons. Journal of neuroscience, 22(8), 3005–3015.

51. Lalanne, T., Oyrer, J., Farrant, M., & Sjöström, P. J. (2018). Synapse type-dependent expression of calcium-permeable AMPA receptors. Frontiers in synaptic neuroscience, 10, 34.

52. McBain, C. J., & Dingledine, (1993). Heterogeneity of synaptic glutamate receptors on CA3 stratum radiatum interneurones of rat hippocampus. The Journal of physiology, 462(1), 373–392.

53. McMoneagle, E., Zhou, J., Zhang, S., Huang, W., Josiah, S. S., Ding, K., … & Zhang, J. (2024). Neuronal K+-Cl-cotransporter KCC2 as a promising drug target for epilepsy treatment. Acta Pharmacologica Sinica, 45(1), 1–22.

54. Mellor, I. R., Brier, T. J., Pluteanu, F., Strømgaard, K., Saghyan, A., Eldursi, N., … & Usherwood, P. N. R. (2003). Modification of the philanthotoxin-343 polyamine moiety results in different structure-activity profiles at muscle nicotinic ACh, NMDA and AMPA receptors. Neuropharmacology, 44(1), 70–80.

55. Miles, R., Blaesse, P., Huberfeld, G., Wittner, L., & Kaila, K. (2012). Chloride homeostasis and GABA signaling in temporal lobe epilepsy. Jasper’s Basic Mechanisms of the Epilepsies [Internet]*. 4th edition*.

56. Miya, K., Takahashi, Y., & Mori, H. (2014). Anti-NMDAR autoimmune encephalitis. Brain and Development, 36(8), 645–652.

57. Mony, L., Zhu, S., Carvalho, S., & Paoletti, P. (2011). Molecular basis of positive allosteric modulation of GluN2B NMDA receptors by polyamines. The EMBO journal, 30(15), 3134–3146.

58. Peng, Y., Bjelde, A., Aceituno, P. V., Mittermaier, F. X., Planert, H., Grosser, S., … & Geiger, J. R. (2024). Directed and acyclic synaptic connectivity in the human layer 2-3 cortical microcircuit. Science, 384(6693), 338–343.

59. Plested, A. J., & Poulsen, M. H. (2021). Crosslinking glutamate receptor ion channels. In Methods in Enzymology (Vol. 652, pp. 161-192). Academic Press.

60. Riva, I. (2020). Biophysical properties of AMPA receptor complexes. Humboldt Universitaet zu Berlin (Germany).

61. Rogawski, M. A. (1992). The NMDA receptor, NMDA antagonists and epilepsy therapy: a status report. Drugs, 44(3), 279–292.

62. Rogawski, M. A. (2013). AMPA receptors as a molecular target in epilepsy therapy. Acta Neurologica Scandinavica, 127, 9–18.

63. Sadeghi, M. A., Hemmati, S., Mohammadi, S., Yousefi-Manesh, H., Vafaei, A., Zare, M., & Dehpour, A. R. (2021). Chronically altered NMDAR signaling in epilepsy mediates comorbid depression. Acta Neuropathologica Communications, 9(1), 53.

64. Salpietro, V., Dixon, C. L., Guo, H., Bello, O. D., Vandrovcova, J., Efthymiou, S., … & Houlden, H. (2019). AMPA receptor GluA2 subunit defects are a cause of neurodevelopmental disorders. Nature communications, 10(1), 3094.

65. Sanderson, J. L., Gorski, J. A., & Dell’Acqua, M. L. (2016). NMDA receptor-dependent LTD requires transient synaptic incorporation of Ca2+-permeable AMPARs mediated by AKAP150-anchored PKA and calcineurin. Neuron, 89(5), 1000–1015.

66. Schwarz, N., Hedrich, U. B., Schwarz, H., PA, H., Dammeier, N., Auffenberg, E., … & Koch, H. (2017). Human Cerebrospinal fluid promotes long-term neuronal viability and network function in human neocortical organotypic brain slice cultures. Scientific reports, 7(1), 12249.

67. Sommer, B., Köhler, M., Sprengel, R., & Seeburg, P. H. (1991). RNA editing in brain controls a determinant of ion flow in glutamate-gated channels. Cell, 67(1), 11–19.

68. Shepherd, J. D., & Huganir, R. L. (2007). The cell biology of synaptic plasticity: AMPA receptor trafficking. Annu. Rev. Cell Dev. Biol., 23(1), 613–643.

69. Sobolevsky, A. I., Rosconi, M. P., & Gouaux, E. (2009). X-ray structure of AMPA-subtype glutamate receptor: symmetry and mechanism. Nature, 462(7274), 745.

70. Sommer, B., Köhler, M., Sprengel, R., & Seeburg, P. H. (1991). RNA editing in brain controls a determinant of ion flow in glutamate-gated channels. Cell, 67(1), 11–19.

71. Tóth, K., & McBain, C. J. (1998). Afferent-specific innervation of two distinct AMPA receptor subtypes on single hippocampal interneurons. Nature neuroscience, 1(7), 572–578.

72. Traynelis, S. F., Wollmuth, L. P., McBain, C. J., Menniti, F. S., Vance, K. M., Ogden, K. K., … & Dingledine, R. (2010). Glutamate receptor ion channels: structure, regulation, and function. Pharmacological reviews, 62(3), 405–496.

73. Trott, O., & Olson, A. J. (2010). AutoDock Vina: improving the speed and accuracy of docking with a new scoring function, efficient optimization, and multithreading. Journal of computational chemistry, 31(2), 455–461.

74. Twomey, E. C., Yelshanskaya, M. V., Vassilevski, A. A., & Sobolevsky, A. I. (2018). Mechanisms of channel block in calcium-permeable AMPA receptors. Neuron, 99(5), 956–968.

75. van van Hugte, E. J., Schubert, D., & Nadif Kasri, N. (2023). Excitatory/inhibitory balance in epilepsies and neurodevelopmental disorders: Depolarizing γ-aminobutyric acid as a common mechanism. Epilepsia, 64(8), 1975–1990.

76. Vollmar, W., Gloger, J., Berger, E., Kortenbruck, G., Köhling, R., Speckmann, E. J., & Musshoff, U. (2004). RNA editing (R/G site) and flip–flop splicing of the AMPA receptor subunit GluR2 in nervous tissue of epilepsy patients. Neurobiology of disease, 15(2), 371–379.

77. Wakazono, Y., Midorikawa, R., & Takamiya, K. (2024). Temporal and quantitative analysis of the functional expression of Ca2+-permeable AMPA receptors during LTP. Neuroscience Research, 198, 21–29.

78. Wahab, A., Albus, K., Gabriel, S., & Heinemann, U. (2010). In search of models of pharmacoresistant epilepsy. Epilepsia, 51, 154–159.

79. Wang, T., Wang, J., Cottrell, J. E., & Kass, I. S. (2004). Small physiologic changes in calcium and magnesium alter excitability and burst firing of CA1 pyramidal cells in rat hippocampal slices. Journal of neurosurgical anesthesiology, 16(3), 201–209.

80. Washburn, M. S., & Dingledine, R. (1996). Block of alpha-amino-3-hydroxy-5-methyl-4-isoxazolepropionic acid (AMPA) receptors by polyamines and polyamine toxins. Journal of Pharmacology and Experimental Therapeutics, 278(2), 669–678.

81. Washburn, M. S., Numberger, M., Zhang, S., & Dingledine, R. (1997). Differential dependence on GluR2 expression of three characteristic features of AMPA receptors. Journal of Neuroscience, 17(24), 9393–9406.

82. Wilbers, R., Galakhova, A. A., Driessens, S. L., Heistek, T. S., Metodieva, V. D., Hagemann, J., … & Goriounova, N. A. (2023). Structural and functional specializations of human fast-spiking neurons support fast cortical signaling. Science advances, 9(41), eadf0708.

83. Wood, J. N., Bevan, S. J., Coote, P. R., Dunn, P. M., Harmar, A., Hogan, P., … & Wheatley, S. (1990). Novel cell lines display properties of nociceptive sensory neurons. Proceedings of the Royal Society of London. Series B: Biological Sciences, 241(1302), 187–194.

84. Zhao, Y., Chen, S., Swensen, A. C., Qian, W. J., & Gouaux, E. (2019). Architecture and subunit arrangement of native AMPA receptors elucidated by cryo-EM. Science, 364(6438), 355–362.

85. Zinchenko, V. P., Teplov, I. Y., Kosenkov, A. M., Gaidin, S. G., Kairat, B. K., & Tuleukhanov, S. T. (2024). Participation of calcium-permeable AMPA receptors in the regulation of epileptiform activity of hippocampal neurons. Frontiers in synaptic neuroscience, 16, 1349984.

