## Supplementary Figure 1 for "The polyamine naphthyl-acetyl spermine trihydrochloride (NASPM) lacks specificity for Ca^2+^-permeable AMPA receptors and suppresses seizure like activity in human brain tissue by inhibition of NMDA receptors"

# S1

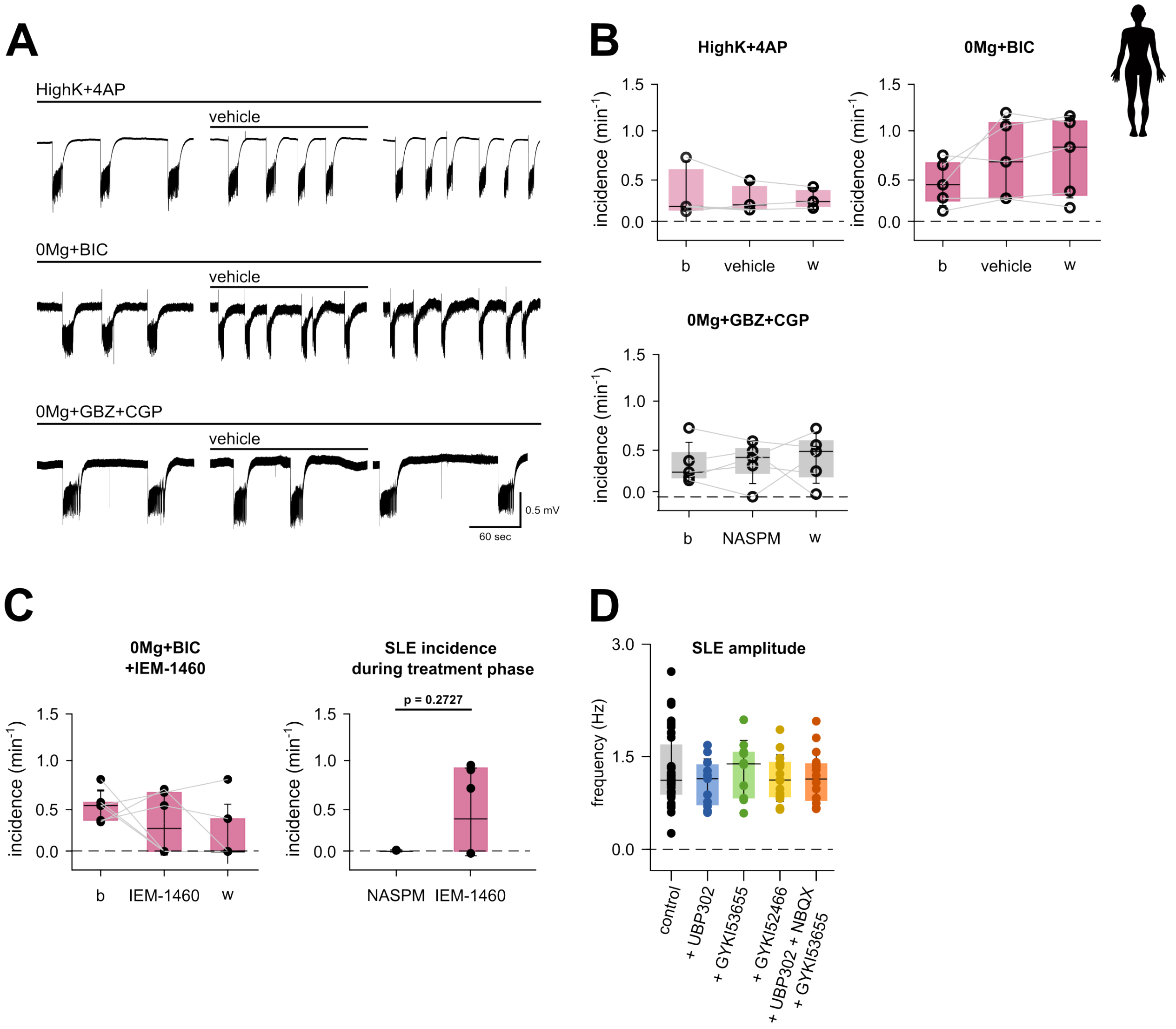


**Supplementary Figure 1. Vehicle or IEM-1460 intervention does not affect SLEs incidence. SLE amplitude and duration is not affected by glutamatergic blockers UBP302, GYKI53655, GYKI5245, NBQX.**

**A.** Vehicle intervention does not significantly reduce SLEs incidence during intervention phase. For each induction method a representative portion of baseline, vehicle intervention and washout phase are shown, slow drift of the traces were removed for better visualization **B.** SLE incidence remains stable over time during the intervention of 300 µL Milli-Q water as vehicle in all induction methods studied (High+4AP, 5 slices from 3 patients; 0Mg+BIC, 7 slices from 5 patients; 0Mg+GBZ+CGP, 6 slices from 5 patients). Statistics**:** Friedman test with Dunn's multiple comparisons; normality was assessed via Shapiro-Wilk test. **C.** Similarly, IEM-1460 intervention does not affect significantly SLEs incidence in a 0Mg+BIC induction method (left panel). Statistics**:** Friedman test with Dunn's multiple comparisons; normality assessed via Shapiro-Wilk test. A significant difference in observed in SLE incidence during intervention of 100 µM NASPM (9 slices from 5 patients) or 100µM IEM-1460 (9 slices from 6 patients, $p = 0.2727$, right panel). Statistics**:** Mann-Whitney test; normality was assessed via Shapiro-Wilk test. All data are shown as scatter plots of individual recordings; superimposed box plots indicate the median (centre line), interquartile range (box edges), and standard deviation (whiskers). **D.** In the induction method 0Mg+GBZ+CGP with addition of various glutamatergic UBP302, GYKI53655, GYKI52455, and a combination of UBP302, GYKI53655, and NBQX, SLE amplitude and duration doesn´t vary significantly across the methods. 0Mg+GBZ+CGP+

NASPM and +APV as intervention compounds are shown for completeness. Statistics: Kruskal-Wallis with Dunn's multiple comparisons. Normality assessed via Shapiro-Wilk test. Sample sizes: 0Mg+GBZ+CGP (n=40 slices from 16 patients); +UBP302 (n=11 slices from 7 patients); +GYKI53655 (n=11 slices from 7 patients); +GYKI52466 (n=19 slices from 7 patients); +UBP302+GYKI53655+NBQX (n=22 slices from 10 patients).
